# Topsicle: a method for estimating telomere length from whole genome long-read sequencing data

**DOI:** 10.1101/2025.07.10.664126

**Authors:** Linh Nguyen, Jae Young Choi

## Abstract

Telomeres protect chromosome ends and its length varies significantly between organisms. Because telomere length variation is associated with various biomedical and eco-evolutionary phenotypes, many biological fields are interest in understanding its biological significance. Here we introduce Topsicle, a computational method that estimates telomere length from whole genome long read sequencing data using k-mer and change point detection analysis. Simulations showed Topsicle was robust to sequencing errors and coverage. Application of Topsicle on plant and human cancer cells showed high accuracy and comparable results to direct telomere length measurements. We predict Topsicle will be a useful tool for studying telomere biology.

## Background

Telomeres are nucleoprotein complexes found at the ends of eukaryotic chromosomes and protect the genome against instability [1]. During DNA replication the lagging strand of chromosome ends will lose DNA with every cellular replication (*i.e.* the chromosome end replication problem [2,3]). In addition, exposed DNA from the telomere has the potential to be mistaken as damaged DNA resulting in cellular arrest (*i.e.* the chromosome end protection problem [4]). To prevent such issues, telomeres are bound by specialized proteins for protection and the telomerase maintains chromosomal ends through reverse transcribing telomere DNA sequences from a noncoding RNA gene [5,6]. Abnormal telomere length can have catastrophic consequences, where shortened telomeres can result in chromosome end-to-end fusions [7,8] and cellular senescence [9], meanwhile elongated telomeres are typically associated with cancer and tumorigenesis [10,11]. With such deleterious consequences, selection should maintain an optimal telomere length for all organisms.

Telomere length, however, varies substantially between species (*e.g.* telomeres of mice can be 10 times longer than humans [12]), and even within species between individuals the length of the telomere varies significantly as well (*e.g.* human telomere lengths range from 5 to 15 kb [13] and *Arabidopsis thaliana* telomere lengths range from 1 to 12 kbp [14]). Why does the length of a crucial chromosomal structure vary within organisms across multiple kingdoms? The answer might lie in the association between telomere length and life history traits (*i.e.* phenotypes that affect the reproduction and survival of an organism at different developmental stages), which affects fitness (*i.e.* traits that determine reproductive success) and is subjected to natural selection. Because of the association between the telomere and cellular senescence [15,16], hypotheses have proposed telomere length variation is a result of life history tradeoffs relating to the aging trajectory of an organism [17,18]. In mammals the shortening rate of the telomere is correlated with the lifespan of different species [19], consistent with the model that selection on aging underlies the telomere length variation. In plants, on the other hand, the process of aging differs significantly from animals [20] that it has been unclear whether aging or lifespan related traits were also driving their telomere length variation. Recent studies, however, have shown that telomere length is associated with flowering times in three evolutionary divergent plant species *A. thaliana*, maize, and rice [14] and direct manipulation of telomere lengths through mutations in key telomere maintenance genes can alter both vegetative and reproductive traits in *A. thaliana* [21]. In the end, these studies have highlighted the potential biological link between life history strategies and chromosomal structures such as the telomere, but what genetic and evolutionary mechanism underlies this link is largely unknown.

An initial step towards understanding the biological significance of the telomere length variation requires an accurate measurement of the telomere in multiple different individuals, genotypes, or species. The gold standard for measuring telomere length is the terminal restriction fragment (TRF) assay [22], which involves restriction enzymes to digest the DNA to release the telomeric region and visualized using a Southern blot with a telomere sequence probe [23]. TRF is a highly accurate method for measuring the length of the telomere, but the method does have challenges and drawbacks that acts as a barrier for widespread usage in a standard laboratory, which includes the requirement of high amounts of DNA (∼5 μg), the requirement of radioactive primers (although see [24] for non-radioactive based labeling) that are not easily accessible, and depending on the restriction enzyme it could incorporate subtelomeric regions into the length estimates. Alternative methods have combined microscopy and chromosomal labeling such as quantitative fluorescence in situ hybridization (Q-FISH) that quantifies telomere length signals from cells with metaphase chromosomes [25], and Flow-FISH that uses flow cytometry and measures telomere length from individual cells [26]. These microscopy based methods require cells from a specific developmental stage, which may not be readily available depending on the organism. In addition, both TRF and microscopy based methods are labor intensive and it is a challenge to scale up and conduct high throughput screens in many samples. On the other hand, increased throughput has been accomplished by applying quantitative PCR (qPCR) and quantifying the amount of telomere repeats as a way to approximate telomere lengths [27]. But due to the inherent complications of amplifying a tandemly repeating sequence such as the telomere, a qPCR based method requires careful designing of PCR primers to avoid dimerization and careful calibration to account for potential differences in PCR conditions between samples. In this regard, noticeable variation in qPCR based telomere length results between laboratories has raised concerns of adopting the qPCR method for length measurements [28]. Lastly, none of the aforementioned techniques are designed to estimate telomere lengths of specific chromosomes. To measure telomere lengths from specific chromosomes, methods such as STELA [29] and PETRA [30] were developed but these require known subtelomeric sequences to design primers and amplify the telomeres of specific chromosomes. We refer to [31] for an in depth review on the advantages and disadvantages of each method.

With major advancements in genome sequencing technology, computationally based methods have become an alternative approach for estimating telomere lengths. Methods such as TelSeq [32], K-seek [33,34], Computel [35], Telomerecat [36], and TelomereHunter [37] have been used to analyze short read sequencing data (usually generated from the Illumina sequencing platform) for estimating the telomere length of a sample using an approach analogous to a qPCR based method. These short read sequencing based bioinformatic methods have a common theme of quantifying the telomere repeat in every sequencing read and the total abundance of the repeat represents the sample’s telomere length. But before comparing telomere length estimates between samples, proper normalizations should be done to reduce batch effects between sequencing libraries [38]. For instance, a tagmentation based or physically shearing based fragmentation of DNA can result in biases in the sequenced genomic regions [39], or PCR amplification on DNA fragments after adapter ligation could introduce amplification bias during sequencing [40,41]. Overall, short read sequencing based computational methods are well correlated with experimentally based methods. For example, across 112 A. thaliana ecotypes Spearman’s *ρ* is 0.55 between K-seek versus TRF based telomere length estimates [14], and across 100 Populus genotypes Peasrson’s *r* is 0.62 between K-seek versus q-PCR based telomere length estimates [42]. A major advantage of computationally based methods is the ease of estimating telomere length as it only requires a whole genome sequencing dataset. Hence computationally based approaches can be a powerful method for investigating telomere length variation.

Long read sequencing technology (advanced by Pacific Biosciences (PacBio) and Oxford Nanopore Technologies (Nanopore)) are revolutionizing the genomics field [43] and it has major potential to be a powerful computational tool for investigating the telomere length variation within population and between species. Read length from long read sequencing platforms are orders of magnitude longer than short read sequencing platforms (tens of kilobase pairs versus 100–300 bp). These long reads have greatly aided in resolving the complex and highly repetitive regions of the genome [44], and near gapless genome assemblies (also known as telomere-to-telomere assembly) are generated for multiple organisms [45,46]. The long read sequences can also be used for estimating telomere length, since whole genome sequencing using a long read sequencing platform would contain reads that span the entire telomere and subtelomere region. Computational methods can then be developed to determine the telomere– subtelomere boundary and use it to estimate the telomere length. As an example, telomere-to-telomere assemblies have been used for estimating telomere length by analyzing the sequences at the start and end of the gapless chromosome assembly [47–50]. But generating gapless genome assemblies are resource intensive and can’t be used for estimating the telomeres of multiple individuals. Alternatively, methods such as TLD [51], Telogator [52], and TeloNum [53] analyze raw long read sequences to estimate telomere lengths. These methods require a known telomere repeat sequence but this can be determined through k-mer based analysis [54]. Specialized methods have also been developed to concentrate long reads originating from chromosome ends. These methods involve attaching sequencing adapters that are complementary to the single-stranded 3’ G-overhang of the telomere, which can subsequently be used for selectively amplifying the chromosome ends for long read sequencing [55–58]. While these methods can enrich telomeric long reads, it requires optimization of the protocol (e.g. designing the adapter sequence to target the G-overhang) and organisms with naturally blunt-ended telomeres [59,60] would have difficulty implementing the methods.

An explosion of long read sequencing data has been generated for many organisms across the animal and plant kingdom [61,62]. A computational method that can use this abundant long read sequencing data and estimate telomere length with minimal requirement can be a powerful toolkit for investigating the biology of telomere length variation. But so far such a method is not available and implementing one would require addressing two major algorithmic considerations before it can be widely used across many different organisms. The first algorithmic consideration is the ability to analyze the diverse telomere sequence variation across the tree of life. All vertebrates have an identical telomere repeat motif TTAGGG [63] and most previous long read sequencing based computational methods were largely designed for analyzing human genomic datasets where the algorithms are optimized on the TTAGGG telomere motif. But the telomere repeat motif is highly diverse across the animal and plant kingdom [64–67], and there are even species in fungi and plants that utilize a mix of repeat motif resulting in a sequence complex telomere structure [64,68,69]. A new computational method would need to accommodate the diverse telomere repeat motif, especially across the inherently noisy and error prone long read sequencing data [70]. With recent improvements in sequencing chemistry and technology (HiFi sequencing for PacBio and Q20+ Chemistry kit for Nanopore) error rates have been substantially reduced to <1% [71,72]. But even with this low error rate a telomeric region that is several kilobase pairs long can harbor substantial erroneous sequences across the read [73] and hinder the identification of the correct telomere–subtelomere boundary. In addition, long read sequencers are especially error prone to repetitive homopolymer sequences [74–76], and the GT rich microsatellite telomere sequences are predicted to be an especially erroneous region for long read sequencing. A second algorithmic consideration relates to identifying the telomere– subtelomere boundary. Prior long read sequencing based methods [51,52] have used sliding windows to calculate summary statistics and a threshold to determine the boundary between the telomere and subtelomere. Sliding window and threshold based analyses are commonly used in genome analysis, but it places the burden on the user to determine the appropriate cutoff, which for telomere length measuring computational methods may differ depending on the sequenced organism. In addition, threshold based sliding window scans can inflate both false positive and false negative results [77–82] if the cutoff is improperly determined.

Here we introduce Topsicle, a computational method that uses a novel strategy to estimate telomere lengths from raw long read sequences from the entire whole genome sequencing library. Methodologically, Topsicle iterates through different substring sizes of the telomere repeat sequence (i.e. telomere k-mer) and different phases of the telomere k-mer are used to summarize the telomere repeat content of each sequencing read. The k-mer based summary statistics of telomere repeats are then used for selecting long reads originating from telomeric regions. Topsicle uses those putative reads from the telomere region to estimate the telomere length by determining the telomere–subtelomere boundary through a binary segmentation change point detection analysis [83]. We demonstrate the high accuracy of Topsicle through simulations and apply our new method on long read sequencing datasets from three evolutionarily diverse plant species (*A. thaliana*, maize, and *Mimulus*) and human cancer cell lines. We believe using Topsicle will enable high-resolution explorations of telomere length for more species and achieve a broad understanding of the genetics and evolution underlying telomere length variation.

## Results

### Spatial distribution of telomere repeats in raw long read sequencing data from chromosome ends

Initially, we investigated the distribution of telomere repeats in raw long read sequences originating from chromosome ends and focusing specifically on the reads that were spanning the telomere–subtelomere boundary region. We focused on *A. thaliana* ecotype Col-0 due to the wealth of telomere related knowledge gained from molecular studies [84], and the abundant genomic data that can be used for computational analysis. We obtained a published nanopore sequencing data for Col-0 [85] and aligned the long reads to the *A. thaliana* Col-0 genome assembly generated from long read sequencing [86], which has 8 out of the total 10 chromosome ends completely assembled. Chromosome 2L and 4L telomeres are directly adjacent to the nucleolar organization regions [87] and because of that the assemblies of these regions are incomplete. We first extracted reads aligning to ends of chromosome 1 to examine the spatial distribution of the telomere repeat in raw nanopore sequencing reads. In Fig 1A we display the long reads and highlighted the regions that exactly matched the A. thaliana telomere repeat sequence AAACCCT [88] (Note the conventional denotation would be TTTAGGG to indicate the repeat synthesized by the telomerase on the G-rich strand. Here, we consider it as a simple k-mer and denote it as the alphabetically ordered sequence).

**Figure 1.**
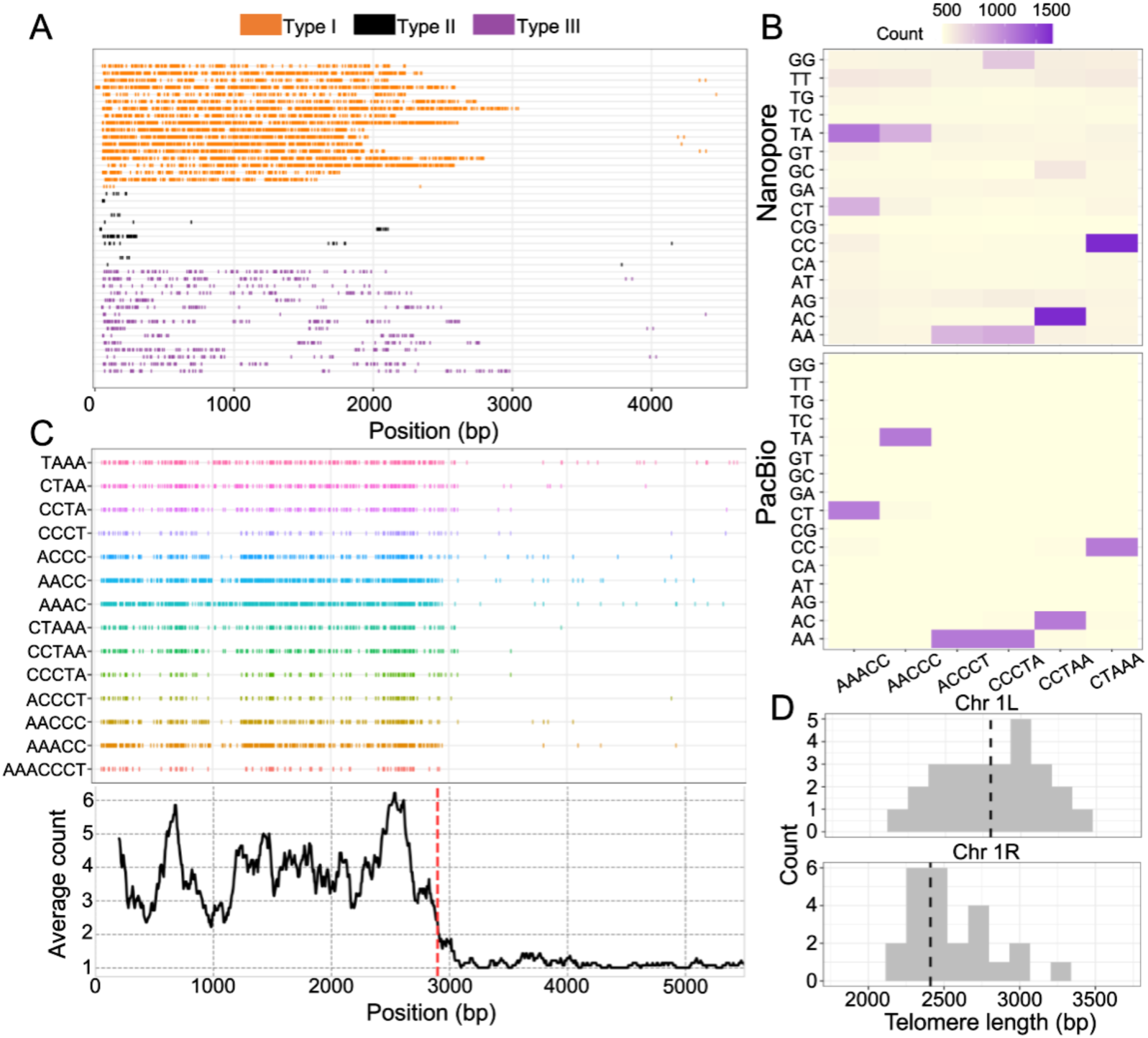
Analysis of telomere repeat k-mers in long read sequences from *A. thaliana* Col-0. (A) Example Type I, II, and III reads from *A. thaliana* Col-0 Nanopore reads. Each color represents an exact match to the telomere repeat AAACCCT and the position found across the sequencing read. Reads are shown so the 5’ end of the sequence is enriched for the AAACCCT repeat and reads with 3’ end enriched for the complement sequence TTTGGGA we show the reverse complement sequence. (B) Co-occurrence heatmap displaying the frequency of a telomere repeat 5-mer (original telomere repeat sequence is AAACCCT) with all possible dinucleotide sequences that can be found at the end of the 5-mer. Top shows frequencies from analyzing reads aligning to chromosome 1R for Nanopore reads and bottom show frequencies from PacBio reads aligning to chromosome 1R. (C) An example Type III Nanopore sequencing read displaying the occurrence of a 4-mer and 5-mer of the original telomere repeat sequence AAACCCT (Top). A sliding window analysis where each window is size 100 bp and it slides 7 bp (bottom). In the window the average k-mer count is calculated and a change point detection method is applied to determine the window where there is a drop in telomere repeat count (red dotted line). (D) Telomere length estimates for Nanopore sequencing reads aligning to chromosome 1L and 1R.

We classified long reads from the telomeric region into three types based on the spatial distribution of the telomere repeat. Type I reads have a high density of AAACCCT sequence at the 5’ region or a high density of TTTGGGA sequence at the 3’ region of the sequencing read. These are likely the long reads spanning the telomere and subtelomere region and the cluster of exact matching telomere repeats suggests these reads have an overall low sequencing errors. A drop in telomere repeat density occurs between 2,000 and 3,000 bp for all reads, which is hypothesized to be where the telomere–subtelomere boundary is located. The telomere length of a read can then be estimated as the length between the start of the telomere repeat and the boundary position. In contrast, Type II reads are those that had minimal telomere repeats across the read. We hypothesized these reads were largely sequenced from the subtelomere region, hence would need to be excluded for downstream analysis as its inclusion would deflate telomere length estimates (our approach of excluding these reads will be discussed later). Type III reads were those that had a cluster of telomere repeats but with large gaps interspersed between clusters. We hypothesized these reads are like Type I reads and originated from the telomere, except there were significant sequencing errors within the telomere regions of the read. Type III can also be used for estimating the telomere length, but the sequencing errors needed to be accommodated in order to accurately determine the telomere–subtelomere boundary for estimating the telomere length of the read.

### Profiling the sequencing errors within the long read sequences from telomeric regions

We investigated the sequencing errors of Type III reads but specifically focusing on the errors involving the telomere repeat sequence. The sequencing errors were examined by quantifying the frequency of the nucleotides that ended after a telomere repeat k-mer. For instance in *A. thaliana*, if there are low sequencing errors an AAACC 5-mer would predominantly end with CT dinucleotides in the long reads. But due to sequencing errors we will also observe non-CT dinucleotides and if the sequencing errors were random we would expect similar frequencies of the non-CT dinucleotides. We investigated this expectation by iterating through all phases of a telomere repeat k-mer (*e.g.* for 5-mer it is AAACC, AACCC, ACCCT, CCCTA, CCTAA, CTAAA, and TAAAC) and counted the frequencies of the terminating nucleotides with the condition that the total length will equal to the telomere repeat size (*e.g.* for 5-mer it will be examining the sequences of the terminating 2 nucleotides).

We used *A. thaliana* Col-0 nanopore reads that aligned to the end of chromosome 1R and investigated its sequencing error by counting the frequency of the dinucleotide sequences terminating the 5-mer telomere repeat. Results showed there were two types of biased sequencing errors for the Nanopore reads (Fig 1B). First involved the CCC homopolymer of the AAACCCT telomere repeat as evidenced by the AAACC 5-mer. For this 5-mer we discovered the most commonly terminating dinucleotide was TA and not CT as expected. This would occur if the AAACCCT sequence was repeatedly mis-basecalled as AAACCT. The second biased sequencing error involved the AA homopolymer as evidenced by the CCCTA 5-mer. We discovered both AA and GG dinucleotides were equally terminating the 5-mer repeats. This indicated the telomere repeat CCCTAAA (Note this is the same sequence as the telomere repeat AAACCCT but in a different phase) was being repeatedly mis-basecalled as CCCTAGG. These results were not due to the sequences in the subtelomere region since an analysis of Col-0 PacBio sequencing reads [89] aligning to chromosome 1R did not show the same frequencies (Fig 1B). For the PacBio sequencing reads, each 5-mer had only a single dinucleotide with high frequency and it corresponded to the expected terminating nucleotide sequence if there were no sequencing errors. In summary, the Nanopore reads had unique biases in the sequencing errors involving the telomere repeat sequence and these would need to be accommodated for an accurate estimation of telomere length.

### Delineating the telomere–subtelomere boundary by summarizing k-mer frequencies

Because of the unique sequencing errors in the Nanopore reads, a search for exact matching telomere repeat sequences would result in the gaps in between telomere clusters as observed for Type III reads (Fig 1A). In addition, the sequencing errors were idiosyncratic (Fig 1B) that it would be hard to predict the error for every Type III read. The erroneous telomere repeats, however, can be detected by analyzing telomere repeat k-mer sequences that were not affected by the sequencing error (i.e. CCTAA or CTAAA from Fig 1B). But because the type of sequencing error would differ for each sequencing read, we quantified the frequency of telomere repeat k-mers of various size and sequence phases across the sequencing read. As an example, we labeled all phases of the 4- and 5-mer with the complete 7-mer telomere repeat for a Type III long read from the *A. thaliana* Col-0 Nanopore sequencing data (Fig 1C). Regions without the 7-mer sequence had matching 4- and 5-mers, suggesting these regions had telomere repeats that were masked by sequencing errors. There was a high density of telomere repeat k-mers up to ∼3,000 bp (Fig 1C), which we hypothesized was the boundary point between the telomere and subtelomere region for the analyzed type III read.

We then investigated if the telomere repeat k-mer density can be used for inferring the telomere length of a sequencing read. The telomere–subtelomere boundary was determined using a sliding window analysis and detecting the window with a significant reduction in the telomere repeat density through change point detection analysis [90]. Change point detection is commonly applied in time series data and aims to determine when there is an abrupt shift in the data series. We applied the binary segmentation method [83] to detect the window with a stepwise change in the telomere repeat k-mer density, and telomere length was measured as the distance between the start of the long read to the telomere– subtelomere boundary position. Applying this to the example Type III read, the estimated telomere length of the read was 2900 bp (Fig 1C).

We further tested our approach by using the Type I and III nanopore reads from chromosome 1 of *A. thaliana* Col-0, and compared it to experimentally based telomere length estimates. In Col-0 Southern blots have shown chromosomes 1L and 1R have similar telomere lengths [30], and previous PETRA analysis from two studies has estimated Col-0 chromosome 1L telomere length as 2,440 bp [21] and nearly 3,000 bp [91]. Long reads aligning to ends of chromosome 1 were manually checked to remove Type II reads, which resulted in 29 reads for chromosome 1L and 31 reads for 1R. For every read we estimated its telomere length and the median telomere length for chromosome 1L was 2800 bp and chromosome 1R was 2410 bp (Fig 1D).

### Telomere Repeat Count (TRC) statistic selects for telomere long reads in A. thaliana

The median telomere length of *A. thaliana* Col-0 chromosome 1 measured from our computational approach was comparable to direct telomere length measurements. But this was conducted on a manually curated dataset of long read sequences that were likely to harbor telomere repeats. We aimed to increase the throughput for selecting the reads containing the telomere repeats by using the same k-mer density approach that was used for measuring telomere length of individual reads. For each sequencing read we calculated a Telomere Repeat Count (TRC) statistic, which represented the density of telomere repeat k-mers at the ends of a sequencing read. Initially, we calculated TRC values using the 4-mer telomere repeat on the manually curated Type I, II, and III reads (TRC distribution for 5-, 6-, and 7-mer repeat are plotted in Figure S1). On the manually curated dataset, the long reads that were visually verified to contain telomere repeats (Type I and III) had significantly higher TRC values (Mann-Whitney U test p-value < 0.0001) compared to the reads that were visually screened to not harbor telomere repeats (Fig 2A). This indicated with an appropriate TRC value cutoff it would be possible to select for long reads that harbored telomere repeats, which can then be used for estimating telomere length. In the case of Type I, II, and III reads that cutoff value may correspond between 0.2 to 0.4.

**Figure 2.**
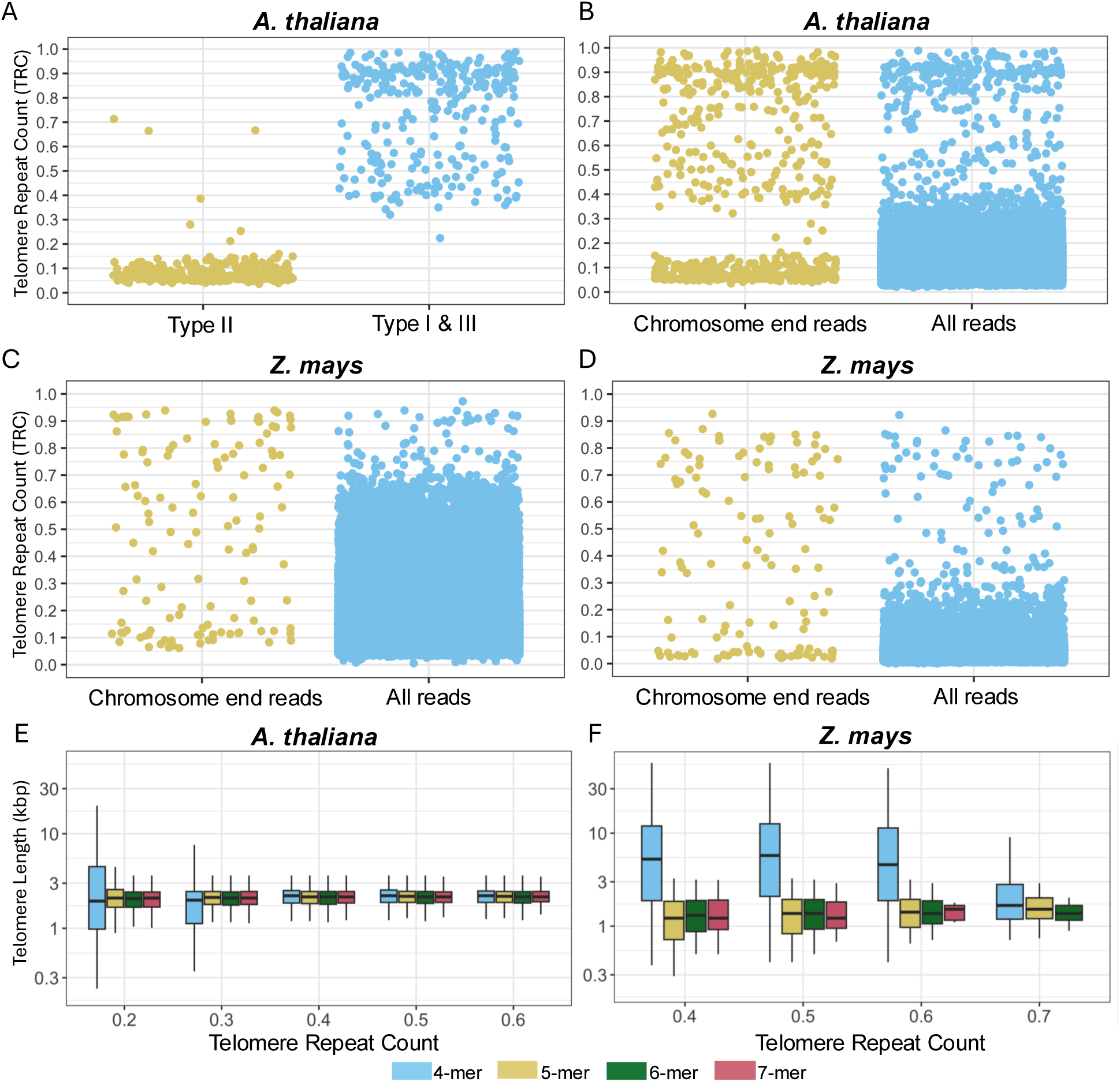
Distribution of Telomere Repeat Count (TRC) values from raw long sequences. (A) TRC values for *A. thaliana* Col-0 Nanopore reads (ERR11436636) that were visually categorized as Type I, II, or III reads. TRC values were calculated using the 4-mers from the telomere repeat sequence AAACCCT. (B) TRC values for *A. thaliana* Col-0 Nanopore reads aligning to chromosome ends or from all sequencing reads. TRC values were calculated using the 4-mers from the telomere repeat sequence AAACCCT. (C) TRC values for maize B73 PacBio reads aligning to chromosome ends or from all sequencing reads. For maize the TRC values were calculated using the reads from a single sequencing library (ERR3288278) out of a total 18 libraries sequenced for B73. TRC values were calculated using the 4-mers from the telomere repeat sequence AAACCCT. (D) TRC values for Maize B73 PacBio reads aligning to chromosome ends or from all sequencing reads. TRC values were calculated using the 5-mers from the telomere repeat sequence AAACCCT. (E) Telomere length estimates from the *A. thaliana* Col-0 Nanopore reads using various TRC value cutoff and k-mer sizes. (F) Telomere length estimates from the maize B73 PacBio reads using various TRC value cutoff and k-mer sizes.

We then investigated an approach that can determine an appropriate TRC cutoff from the whole genome sequencing dataset and select for reads that were likely sequenced from the telomere region. To determine an appropriate TRC cutoff value we first examined the TRC statistics for reads that aligned to chromosome ends and compared it to all long reads from the *A. thaliana* Col-0 Nanopore sequencing library. The former group of reads would be enriched for telomere repeat containing reads (i.e. Type I and III reads) while the latter would be highly enriched for non-telomere repeats (i.e. Type II reads). We analyzed every read that aligned to chromosome ends and calculated TRC values using the frequencies of the 4-mer telomere repeat. From the TRC distribution we could visually identify two groups of reads that were separated by a threshold value of 0.4 (Fig 2B left). We hypothesized the reads with values higher than 0.4 TRC were those sequenced from the telomere region. We then calculated TRC values for all reads from the whole genome sequencing data and results showed a similar distribution as chromosome end reads, where most reads had TRC values lower than 0.4 with a select few with higher than 0.4 TRC values (Fig 2B right). This indicated from the whole genome sequencing data, by selecting for reads with higher than 0.4 TRC values it would selectively choose those that were likely from the telomere region. Median length of the telomere across the selected reads was 2,410 bp (Fig 2E; see boxplot of 4-mer and TRC cutoff of 0.4), which was comparable to the TRF based telomere length for Col-0 at 3,044 bp [14]. We increased the stringency of selecting the long reads that were sequenced from the telomere region by increasing the TRC cutoff (*i.e.* selecting for reads with higher proportion of telomere repeats) and examining longer k-mer sizes (*i.e.* matching closer to the complete telomere repeat sequence), and used the reads that passed this newer conservative criterion to estimate the telomere length. Results showed even at more stringent criteria, the median telomere length estimates did not change and were consistent at ∼2,400 bp (Fig 2E). At lower TRC value cutoff, using the 4-mer telomere repeat resulted in a higher variation in the telomere length estimates but the median was still at ∼2,400 bp.

For Col-0, the ecotype has also been sequenced on a PacBio platform [89] and because we observed differences in error profiles between Nanopore and PacBio reads from the telomere (Fig 1) we investigated the Col-0 telomere length using the PacBio sequencing data. Using the 4-mer telomere repeat and 0.4 TRC value cutoff to search for telomere reads in the PacBio sequencing data, Topsicle calculated the median telomere length as 2,290 bp and this was not significantly different from Nanopore read based estimate (Mann Whitney U test p-value > 0.05 Figure S2). Using the same data we estimated telomere length by searching for a 7-mer (i.e. exact match to AAACCCT telomere repeat) and examined the effect of using a longer k-mer on telomere length estimates. Results showed for PacBio reads the telomere length estimates were not significantly different whether a 4-mer or 7-mer telomere repeat was used for length estimation (Mann Whitney U test p-value = 0.66), but for the Nanopore reads using a 7-mer resulted in significantly shorter telomere length estimates compared to using a 4-mer telomere repeat (Mann Whitney U test p-value = 0.02).

### Application of TRC statistics based filtering on maize long read sequencing data

We then tested if our approach can also be applied to plant species with larger genomes compared to *A. thaliana*. These plant genomes will harbor increased repetitive DNA that could be an obstacle for applying our method. Maize has an identical telomere repeat sequence as *A. thaliana* [92] but its genome size is almost fifteen fold longer (∼2.5 Gb) and more than 80% of its genome consists of repetitive DNA [93,94]. We obtained a PacBio sequencing data for maize genotype B73 [95] and calculated the TRC value for all reads using the 4-mer telomere repeat, which was the parameter also used for *A. thaliana* (Fig 2B). But in contrast to *A. thaliana*, the TRC values for reads that aligned to chromosome ends did not display a bimodal distribution (compare chromosome end read TRC values from Fig 2B and 2C). This indicated in the maize long read sequencing data, there were Type II reads with increased proportions of telomere-like repeats that caused elevated levels of TRC values when the 4-mer repeat is used. We also calculated TRC values for all reads from the whole genome sequencing data using the 4-mer repeat and there was an elevated level of TRC values where majority of reads had TRC value of 0.7 and lower, which contrasts with *A. thaliana* results (compare the TRC values for all reads from Fig 2B and 2C). This suggested compared to analyzing the *A. thaliana* dataset, a more conservative k-mer repeat search or TRC value cutoff would be needed for analyzing the maize dataset.

We increased the stringency of the telomere repeat matching by re-calculating TRC values using the 5-mer telomere repeat (see Figure S3 for 6-mer and 7-mer TRC distributions). Results showed compared to TRC values calculated with the 4-mer telomere repeat, using the 5-mer telomere repeat had fewer reads with high TRC values (Fig 2D). In addition, at a TRC value cutoff between 0.3 to 0.4 the reads from chromosome ends could be divided into two groups where the reads with higher TRC values are likely to be those sequenced from the telomere region. We selected reads that had higher than 0.4 TRC values from the whole genome sequencing data and estimated the telomere length. There were 2,818 reads and the median estimated telomere length was 1,420 bp (Fig 2F; see boxplot of 5-mer and TRC cutoff of 0.4). For B73 the TRF based telomere length estimate was 2,950 bp [92,96]. The estimated telomere length did not change when increased TRC value cutoff or k-mer size were used (Fig 2F), but with the 7-mer (*i.e.* using the complete telomere repeat sequence) there were no telomere long reads with higher than 0.7 TRC values and a telomere length could not be estimated. For the 4-mer telomere repeat, using a TRC value cutoff of 0.7 the telomere lengths were similar to the length estimates from other parameters (Fig 2F). But at lower TRC value cutoffs the telomere lengths were highly elevated, likely due to the inclusion of non-telomere long reads (Fig 2C) during the telomere length estimation.

### Topsicle: a method to estimate telomere length from long sequencing data

From our findings we developed Topsicle (Fig 3), a software package that can analyze long read data generated from whole genome sequencing to search for reads from the telomere region, and use those reads to determine the telomere–subtelomere boundary for estimating the telomere length of the sequenced sample. Topsicle does not require any prior methodology to concentrate telomeric long read sequences as it searches for reads that were likely sequenced from the telomere region by checking for high concentration of telomere repeats using the TRC statistics. An appropriate TRC cutoff to select for telomere long reads can be determined through plotting the TRC distribution for all reads, or by using different incremental cutoff values and investigating the resulting telomere length estimates. For each candidate telomere read, Topsicle estimates the telomere length by quantifying the density of the telomere repeats and demarcating the region of the sequence where there is a significant reduction in the repeat density using change point detection method. Topsicle estimates the telomere length from all reads of a sequencing library and the median length among those reads would represent the overall telomere length of the sequenced sample.

**Figure 3.**
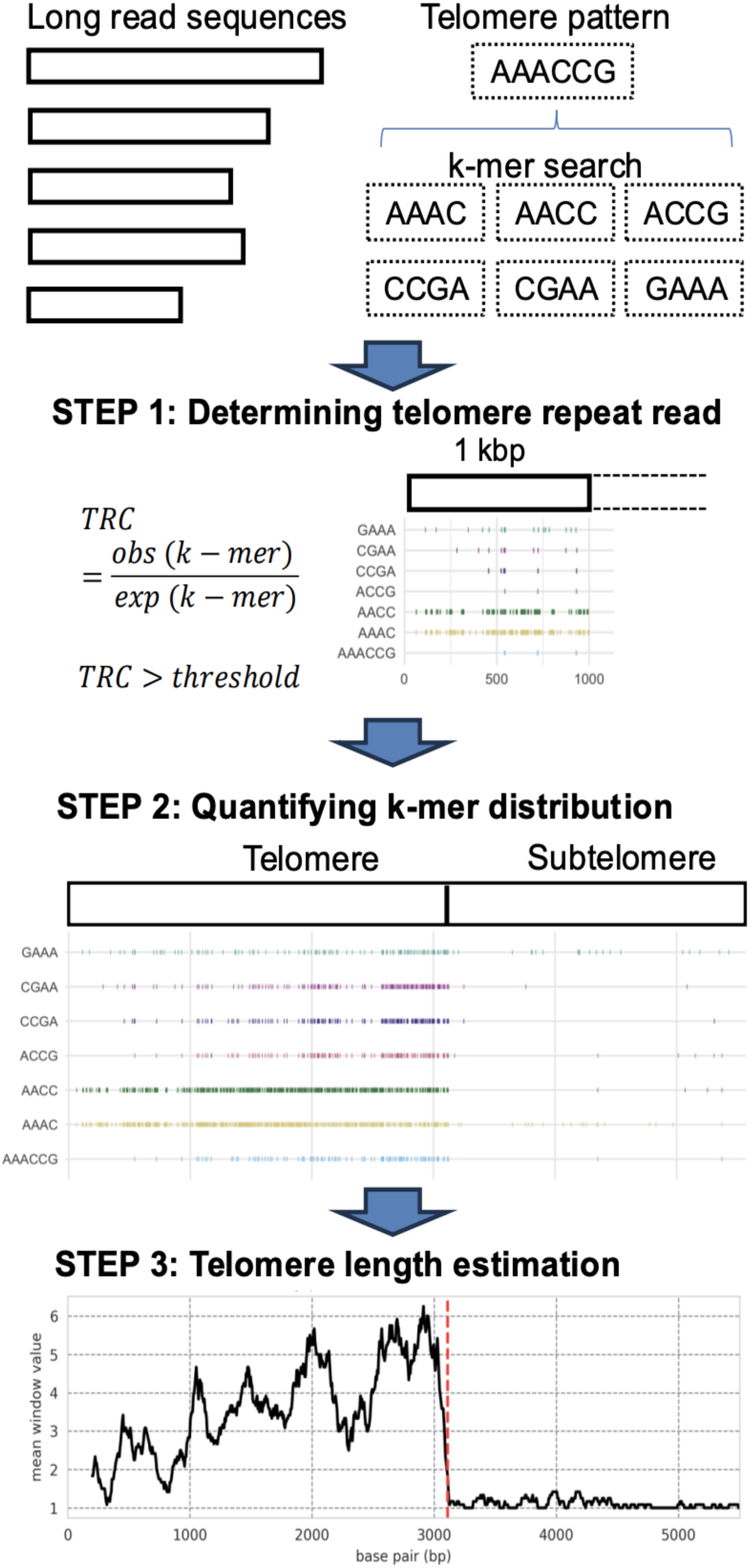
Overview of Topsicle estimating telomere length from long read sequencing data.

### Topsicle can estimate telomere lengths from a reference genome assembly

If a telomere to telomere genome assembly is available, Topsicle can also be used to estimate the length of the telomere directly from the genome assembly. We obtained the reference genomes for *A. thaliana* Col-0, maize B73, and maize Mo17 as they are near telomere to telomere genome assemblies and have TRF based telomere length estimates for comparison. We extracted the chromosome end sequences as inputs for Topsicle and compared it to raw long read based telomere length estimates (Table S1). Results showed the telomere length from the maize B73 reference genome was closer to the TRF based length estimate (difference of 1,950 bp). But for *A. thaliana* Col-0 and maize Mo17 the raw read based estimates were closer to the TRF based length estimates (difference of 634.62 bp and 2409.58 bp respectively). In particular for Mo17, the reference genome based telomere length estimate was over two fold longer than the TRF based length estimates.

### Simulation of Topsicle performance

We investigated the performance of Topsicle through simulations and tested the effects of variations in read length, proportion of the telomere repeats in a read, error rate, and genome-wide coverage can have on the telomere length estimates. Initially, we simulated reads with varying length and proportion of telomere repeat under three different error rates (10%, 20%, and 30% of the sequence) and used Topsicle with the 4-mer telomere repeat search and a TRC value cutoff of 0.4. Results showed Topsicle estimated telomere lengths estimates decreased with increasing sequencing errors but this underestimation was usually <5% of the simulated telomere length (Fig 4A). It is only at very high sequencing error rates of 30% there were more than 10% telomere length underestimation, but this type of errors are not usually seen in recent long read sequencing platforms. Using the same simulated reads we applied longer telomere repeat k-mers and estimated the telomere length. With the 6-mer and 7-mer there were no reads with sequencing error of 30% that passed the 0.4 TRC value cutoff and no telomere length could be estimated. But in lower sequencing errors the size of the k-mer did not largely affect the telomere length estimates and were all <5% of the true telomere length (Figure S4).

**Figure 4.**
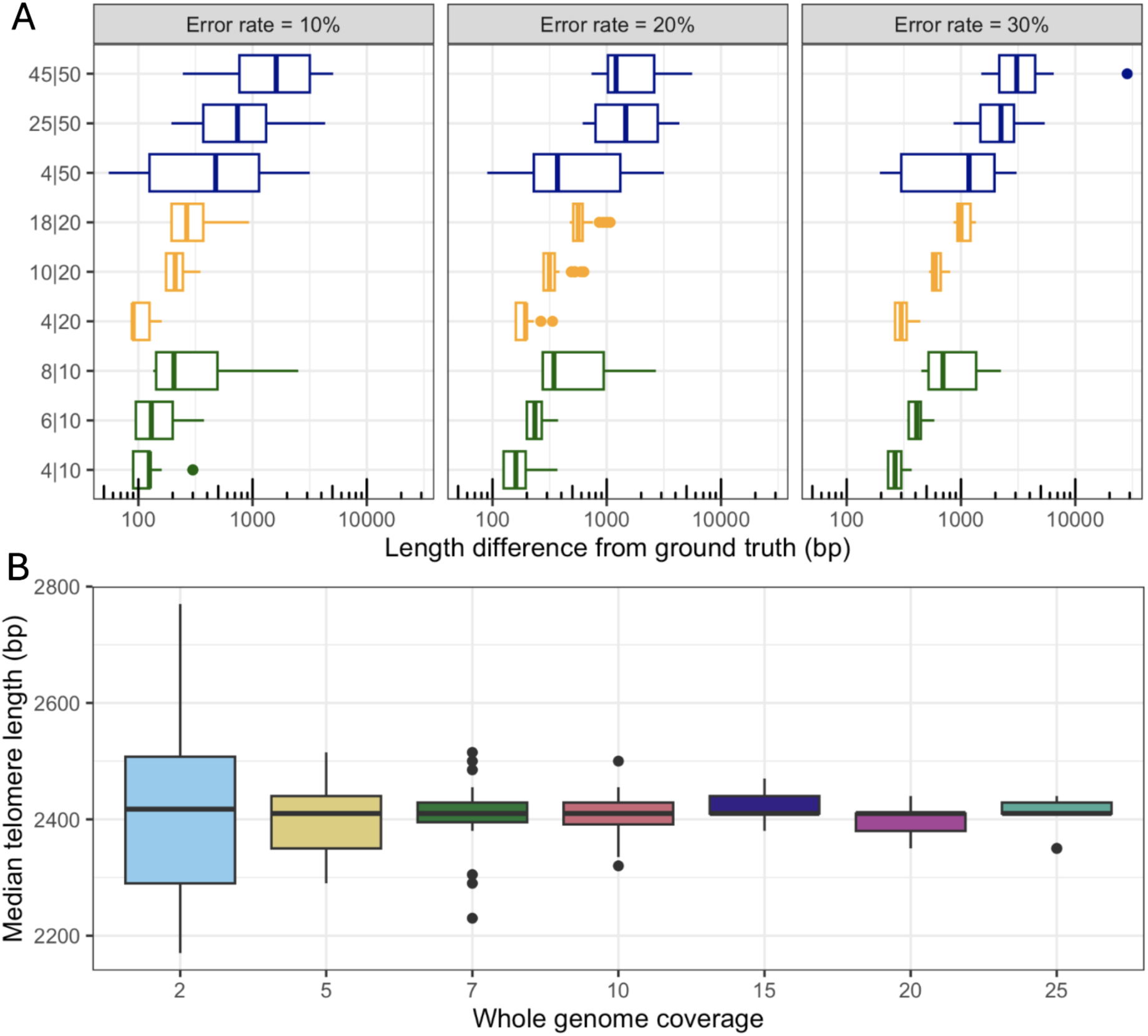
Applying Topsicle on simulated dataset. (A) Telomere length was estimated with Topsicle using the 4-mer and TRC value cutoff of 0.4 on 30 simulated reads with error rates of 10%, 20%, and 30% for reads with varying length and proportion of telomere repeat. For each simulation the read length is indicated on the right side of the bar (“|”) and the length of the telomere repeat is on the left side of the bar. (B) Coverage simulation by random sampling the whole genome sequencing data and using Topsicle to estimate telomere length. For each coverage the random sampling was done 20 times and Topsicle was used to estimate the telomere length using the 4-mer telomere repeat and TRC value cutoff of 0.4 to select for telomere reads. Each point represents the median telomere length from the sampled dataset.

Because recent long read sequencing data have significantly reduced error rates we also simulated reads with lower than 5% error rates and applied Topsicle. We simulated 1%, 2%, and 5% error rates and estimated telomere length using the 4-mer or 7-mer telomere repeat. Results showed under these low error rates, the estimated telomere lengths from Topsicle was within a 100 bp compared to the true telomere length regardless of the telomere length and the sequencing read size (Fig S5).

We next investigated how genome-wide coverage could affect Topsicle estimates of telomere length. Using the *A. thaliana* Col-0 Nanopore sequencing library, we generated datasets that differed in its genome-wide coverage (from 2× to 25×) by randomly subsetting the original dataset, which has 37× genome-wide coverage. Results showed (Fig 4B) at the lowest simulated coverage depth (2×), the median telomere length estimate from each dataset varied the most compared to higher coverage simulations (maximum difference of median of telomere length estimated between two simulated datasets with a coverage of 2× was 600 bp). At higher coverage simulations there was a noticeable reduction in variation across the telomere length estimates between simulated data. In addition, across the simulated coverages the median telomere length estimates were consistent (∼2,400 bp), which is close to the median telomere length estimate when the whole genome sequencing data was used for the telomere length analysis (2,410 bp).

### Applying Topsicle to A. thaliana long read sequencing based population genomic dataset

We applied Topsicle to a long read sequencing based population genomic dataset from two *A. thaliana* studies [85,89] and estimated the telomere length for 104 ecotypes. Results showed our telomere length estimates ranged from 1,720 bp in Ket-10 to 4,720 bp in ws-4 (Table S2), with a mean telomere length of 2,898 ± 608 bp (± standard error) and median telomere length of 2,800 bp (Fig 5A). These telomere lengths were estimated without using the *A. thaliana* reference genome and selecting for reads with TRC values higher than 0.4. We also estimated the telomere length for each ecotype but using reads that were aligning to the chromosome ends of *A. thaliana* reference genome. There was high correlation in the telomere length estimates between reads that were filtered from the whole genome sequencing data and the reads that were aligning to chromosome ends (Figure S6; Pearson’s *r* = 0.93 and p-value < 0.0001; Spearman’s *ρ* = 0.94 and p-value < 0.0001). Further analysis revealed, the reads that were filtered based on the TRC statistic from the whole genome sequencing data were also the same reads that were aligning to chromosome ends (Figure S7), indicating the TRC value based cutoff were selecting for reads that were sequenced from the telomere regions. We investigated if the sequencing library would affect the telomere length estimates for Topsicle, and found no significant correlations between telomere length and number of total sequence, total number of sequenced basepairs, or read length N50 (Figure S8).

**Figure 5.**
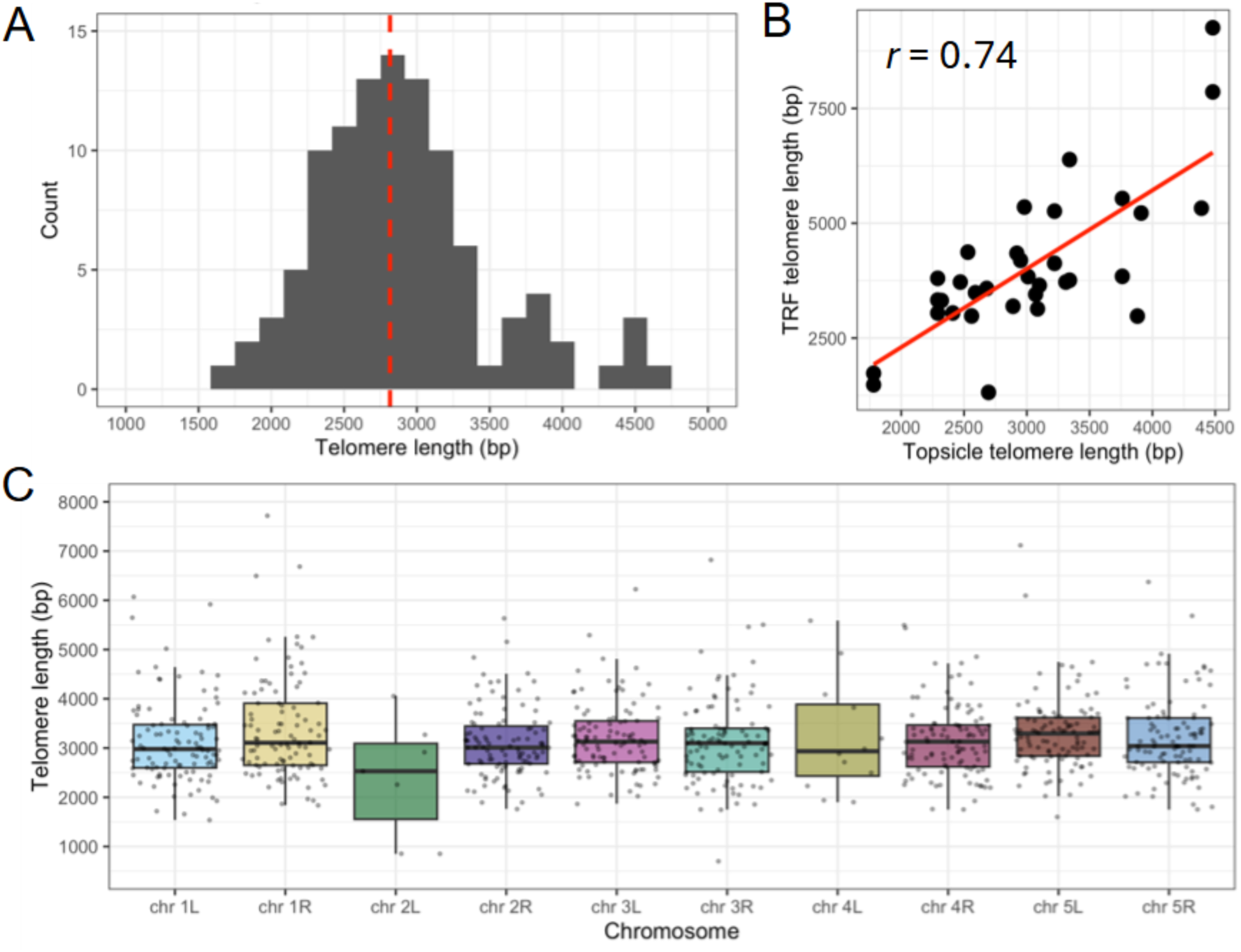
Application of Topsicle on *A. thaliana* long read sequencing dataset. (A) Distribution of telomere length estimates from analyzing whole genome long read sequencing data of 104 *A. thaliana* ecotypes. Red dotted line indicates the median (2815 bp). (B) Scatter plot of telomere lengths from 31 ecotypes that have telomere length estimates from TRF and Topsicle. Pearson’s *r* is shown in the upper-left side and the line of best fit is shown in red line (TRF telomere length = 1.71 × Topsicle telomere length – 1118.13). (C) Boxplot of chromosome specific telomere lengths across the 104 *A. thaliana* ecotypes. Note for chromosome 2L and 4L the majority of the ecotypes did not have reads aligning to those chromosome ends, which prevented the estimation of their chromosome specific telomere length with Topsicle.

For 31 ecotypes there were TRF based telomere length estimates from our previous study [14] and we found a significant positive correlation between the telomere length estimated by Topsicle and TRF (Fig 5B; Pearson’s *r* = 0.74 and p-value < 0.0001; Spearman’s *ρ* = 0.63 and p-value < 0.0001). We noticed the telomere length estimates for Topsicle were lower compared to TRF based estimates, where the median difference was 898.85 bp (Figure S9). We also compared telomere length estimates from Topsicle to telomere repeat abundance calculated by k-Seek, which uses short read Illumina sequencing data. 30 ecotypes had both estimates and there were positive correlations between Topsicle and k-Seek (Figure S10; Pearson’s *r* = 0.559 and p-value < 0.001; Spearman’s *ρ* = 0.301 and p-value = 0.106).

Our telomere length analysis was based on manually choosing a TRC threshold to filter telomere reads, but we explored if an automated procedure could be implemented to choose the appropriate TRC cutoff for length estimation. From Fig 2 we noticed with increasing TRC values the telomere length were converging and at the highest TRC thresholds the telomere length estimates were not changing. Here, an asymptotic TRC point could be determined, which is the threshold point where no further changes in the telomere length are observed with increasing TRC threshold. Using the Lian *et al*. [85] dataset we implemented the asymptotic TRC threshold to calculate the telomere lengths of the long reads and compared it to the hard TRC threshold telomere length estimates from Fig 5A. Results showed high correlation (Pearson’s *r* = 0.99 and p-value < 0.001) between the telomere lengths estimated using a asymptotic TRC threshold and a hard cutoff TRC threshold (Figure S11).

We then took advantage of the chromosome level genome assembly of *A. thaliana* and used the long read population genomic dataset to investigate the chromosome end specific telomere length variation across the ecotypes. A recent study of *A. thaliana* subtelomere sequences adjacent to the telomere revealed high haplotype diversity and genetic structure, where ecotypes had stronger sequence similarities for subtelomeres from the same chromosome ends compared to those from different chromosome ends [97]. This suggested the chromosome end sequences from the reference *A. thaliana* genome can be used for selecting chromosome end specific long reads and use Topsicle to estimate the telomere length. Results showed for each chromosome end (excluding chromosome 2L and 4L), we were able to estimate the telomere length for almost all of the ecotypes (between 96 and 99% of the total ecotypes, Table S3). For chromosome 2L and 4L, because the rDNA cluster is immediately adjacent to the telomere sequence [87], the majority of the ecotypes did not have sequencing reads aligning to the region and a telomere length could not be estimated. But for the rest of the chromosomes, across the ecotypes the Topsicle estimated chromosome end specific telomere lengths had a median length close to 3,000 bp (Fig 5C), and there were no significant differences in length between chromosomes (F_7,810_ =1.393 and p-value = 0.205).

### Applying Topsicle to maize long read sequencing data

We obtained long read sequencing data for 27 maize genotypes [48,95] and applied Topsicle. Because of the large genome size for maize, past studies have sequenced each genotype with multiple libraries ranging from 16 in OH43 to 36 libraries in Mo17 that we analyzed (Table S4). This provided an opportunity to apply Topsicle on each whole genome sequencing library from the same genotype and investigate the telomere lengths from replicated sequencing libraries. We first examined genotype B73 that was sequenced using 18 PacBio libraries and initially analyzed in this study to explore the TRC statistics (Fig 2). For each sequencing library we applied Topsicle using a 5-mer repeat and a TRC cutoff of 0.4 to estimate the library specific telomere length variation. Comparing the lengths between libraries there were no significant differences (Fig 6A; F_17,918_ = 1.49, p-value = 0.09). We then examined genotype OH43, which has a four fold longer telomere compared to B73, and examined the potential variation in telomere length between different sequencing libraries. OH43 was sequenced using 16 libraries that had a median telomere length of 6,705 bp, and there were no significant differences in telomere length between the libraries (Fig 6B; F_15,1113_ = 1.438, p-value = 0.122). For all 27 genotypes, none had significant differences in telomere length between sequencing libraries, highlighting the consistency of Topsicle based length estimates between libraries (Table S5; 5 genotypes had ANOVA p-value < 0.05 but these were not significant after multiple hypothesis correction).

**Figure 6.**
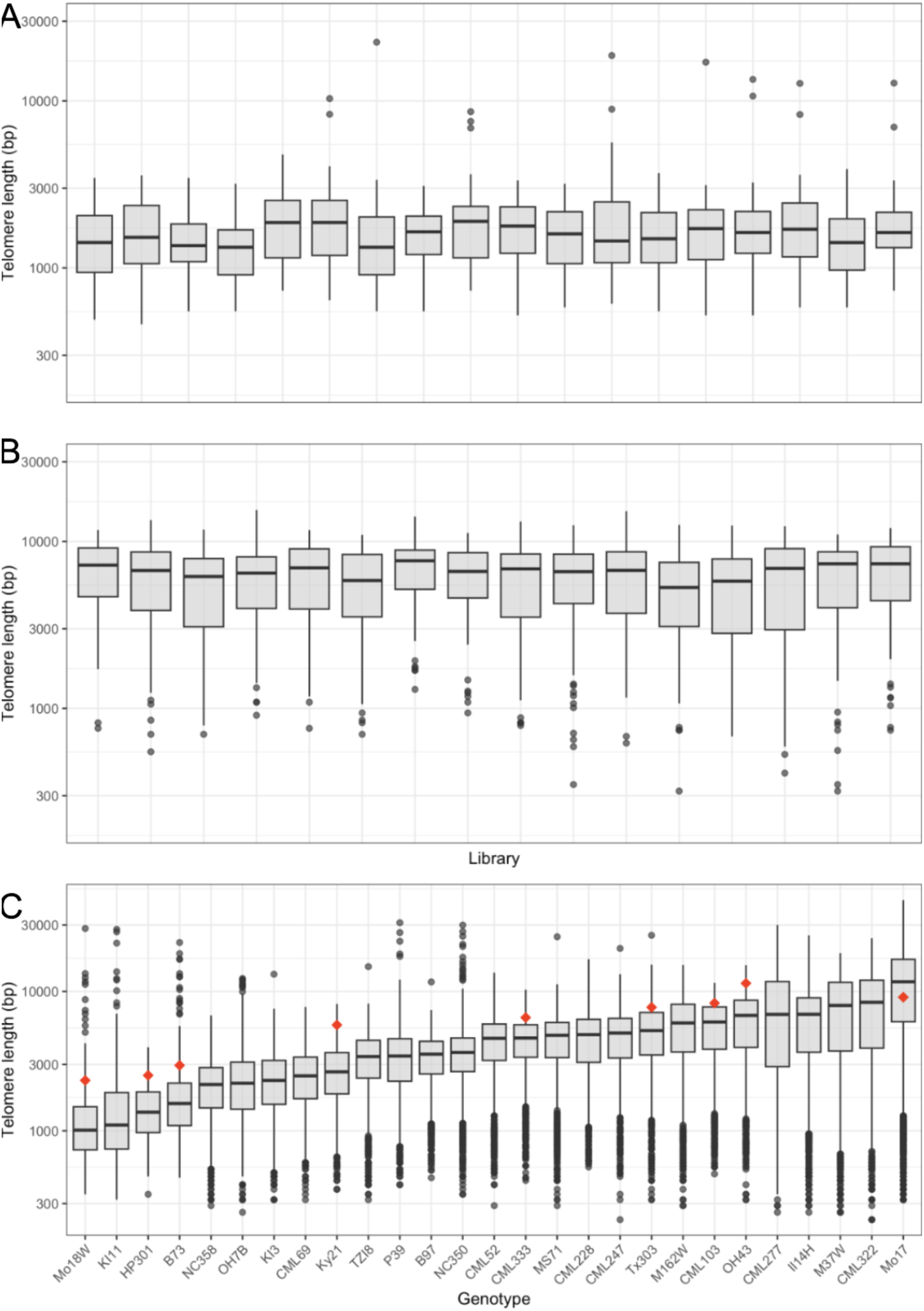
Application of Topsicle on maize long read sequencing dataset. (A) Telomere length estimates for the 18 PacBio libraries for B73. (B) Telomere length estimates for the 16 PacBio libraries for OH43. (C) Boxplot of telomere length estimates for the 27 maize genotypes using Topsicle. The plot shows the telomere length estimates of reads from all sequencing libraries of a genotype that passed a TRC cutoff of 0.4 calculated using a 5-mer repeat sequence. For the genotypes that were previously analyzed with TRF, the telomere length estimates are indicated with a red diamond.

Across the 27 genotypes there was over ten fold variation in telomere lengths (Fig 6C) where the shortest was 1,010 bp in Mo18W and longest was 11,745 bp in Mo17, with a mean telomere length of 5356 ± 25 bp (± standard error) and median telomere length of 4130 bp. These telomere lengths were estimated without using a maize reference genome and selecting for reads with TRC values higher than 0.4 that was calculated using a 5-mer repeat sequence. Nine of the genotypes have been previously assayed with TRF to estimate their telomere lengths [92,96], and Topsicle based telomere length estimates were well correlated with TRF estimates (Fig 6C; Pearson’s *r* = 0.81 and p-value < 0.0001; Spearman’s *ρ* = 0.95 and p-value < 0.0001). But Topsicle estimates were on average 1358.2 bp shorter compared to TRF based telomere length estimates.

Length estimate from TRF will include subtelomere region sequences in addition to the telomere DNA. This could explain why TRF based telomere lengths were longer compared to Topsicle based telomere lengths. To test this we conducted an *in silico* TRF analysis by using the highly contiguous genome assembly of maize genotype B73 [94]. Previously the restriction enzymes Alul, HaeIII and Mbol were used to digest genomic DNA for the TRF protocol and resulting in a telomere length of 3.03 kbp for B73 [96]. We identified the cut site in the subtelomere region of the B73 genome assembly and the *in silico* TRF telomere length was measured as distance between start or end of the chromosome to the cut site. Results showed the median telomere length of the *in silico* TRF was 2.86 kbp which was 1.10 kbp longer than the Topsicle based telomere length (Table S6). This indicated TRF based telomere lengths were longer due to the subtelomere regions being incorporated into the length measurement estimates.

### Applying Topsicle to Mimulus species with sequence heterogeneous telomeres

Recently, we discovered an intriguing case of telomere evolution occurring across three closely related *Mimulus* species (*M. cardinalis*, *M. lewisii*, and *M. verbenaceus*), where the telomere sequence had changed since the last common ancestor (∼2 million years ago) of the three species [68]. *M. cardinalis* and *M. verbenaceus* synthesizes a telomere with a AAACCG repeat sequence, whereas *M. lewisii* has a telomere with a mixed AAACCG and AAACCCG repeat sequence suggesting the telomere sequence was turning over in *M. lewisii*. We analyzed the Nanopore sequencing data for each species [68] to test if Topsicle can also accurately estimate the length of a telomere that has naturally varying sequence variation.

Because our initial analysis of *A. thaliana* Nanopore reads showed significant evidence of sequencing errors (Fig 1B), we initially examined the telomere repeat k-mer profile from the reads aligning to the telomere regions for each *Mimulus* species (Fig 7A). Specifically, for the AAACCG 6-mer telomere sequence we examined the 4-mer telomere sequence and the terminating dinucleotide sequence, and for the AAACCCG 7-mer telomere sequence we examined the 5-mer telomere sequence and the terminating dinucleotide sequence. Results showed the *Mimulus* Nanopore reads from the telomere had different dinucleotide frequencies compared to *A. thaliana* Nanopore reads. Most noticeable was the absence of the mis-basecalling involving the AA and GG dinucleotides observed in *A. thaliana* Nanopore reads, but there were *Mimulus* Nanopore read specific errors such as the AAACCG telomere repeat being mis-basecalled as AAACCA sequence. In *M. lewisii* the k-mer frequencies were able to show the telomere reads had a mix of 6-mer AAACCG and 7-mer AAACCCG repeats. For example, the 4-mer AACC is followed by the dinucleotide CG and the 5-mer AACCC is followed by the dinucleotide GA in *M. lewisii* but not in its sister species *M. cardinalis* or *M. verbenaceus*.

**Figure 7.**
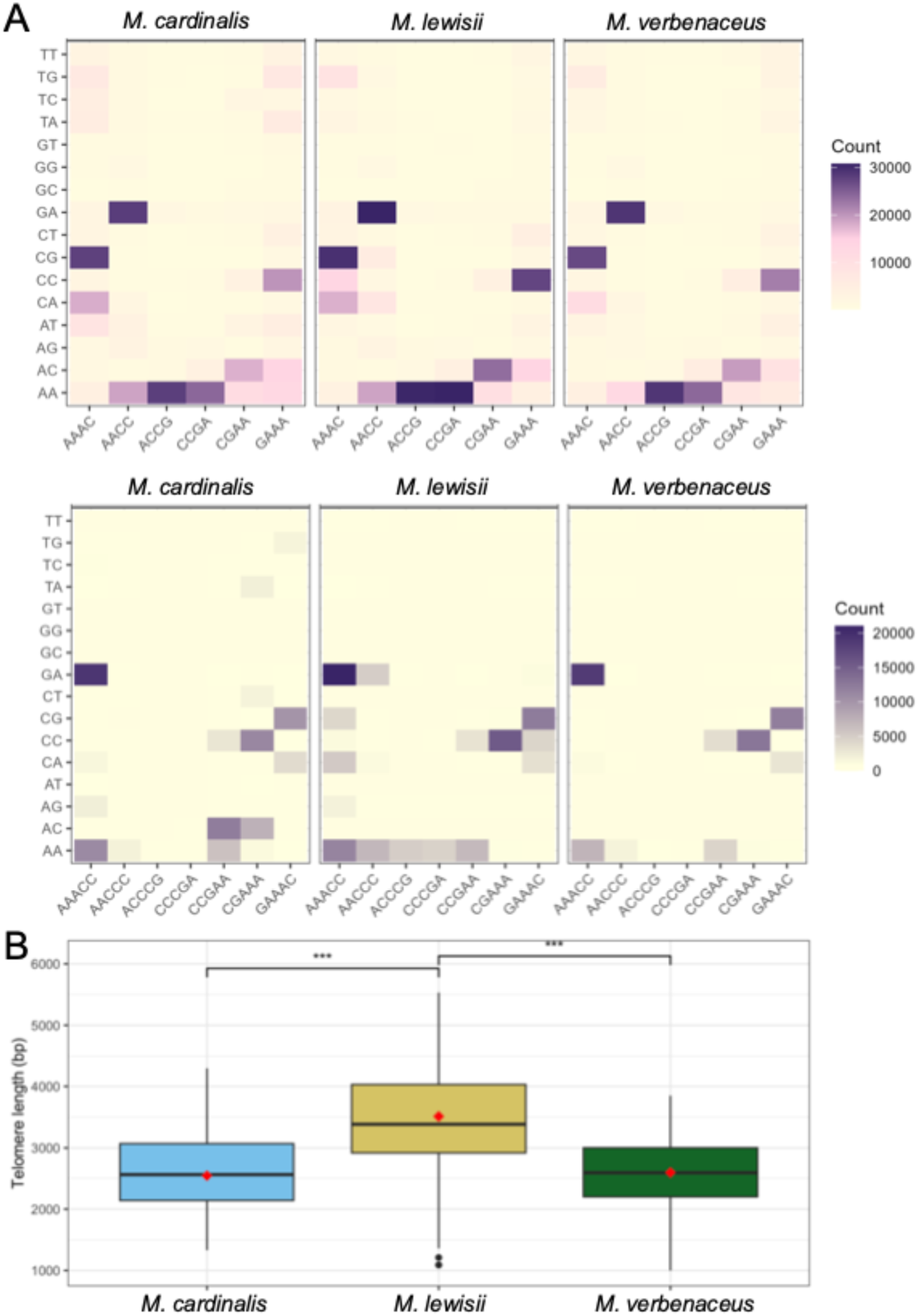
Application of Topsicle on *Mimulus* Nanopore sequencing data. (A) Co-occurrence heatmap displaying the frequency of a telomere repeat. Top shows the telomere repeat 4-mer (original telomere repeat sequence is 6-mer AAACCG) with all possible dinucleotide sequences that can be found at the end of the 4-mer. Bottom shows the telomere repeat 5-mer (original telomere repeat sequence is 7-mer AAACCCG) with all possible dinucleotide sequences that can be found at the end of the 5-mer. Frequencies were counted from Nanopore reads that aligned to chromosome ends of the reference genomes of each respective species. (B) Boxplot of telomere length estimated from Topsicle. For each species the TRF based telomere length is indicated with a red diamond. Significant differences in Topsicle estimated telomere length after a Mann Whitney U test are indicated with *** (p-value < 0.0001).

We estimated the telomere length of each species with Topsicle using the 4-mer of the AAACCG telomere repeat sequence and applying a TRC cutoff of 0.4. The telomere length of *M. cardinalis* was estimated as 2,560 bp, *M. lewisii* as 3,385 bp, and *M. verbenaceus* as 2,590 bp (Fig 7B). These estimates were very similar to the TRF based telomere length estimates of 2,543 bp for *M. cardinalis*, 3,511 bp for *M. lewsii*, and 2,594 bp for *M. verbenaceus* [68]. We then used Topsicle with varying k-mer size (4-mer, 5-mer, and 6-mer of the AAACCG telomere repeat) and TRC cutoff (0.3 to 0.7) and estimated the telomere length for each species. Results showed telomere length estimates were largely consistent across the different parameters. But using an exact match 6-mer AAACCG telomere repeat underestimated the telomere length for M. lewisii at high TRC cutoffs (greater than 0.6), meanwhile for all three species there were no reads that passed a stringent TRC cutoff of 0.7 calculated using a 6-mer (Figure S12).

### Performance of Topsicle on long read sequencing data from human cancer cell lines

Telogator2 [98] is a method that can analyze long read sequencing data from human telomeres and estimate the length of the telomere. We compared the results from Topsicle and Telogator2 by applying each method to the Telo-seq dataset of Schmidt et al. [58], which had enriched for long read sequences from the telomere region and estimated the telomere length of each sequencing read for 10 cancer cell lines. We applied Topsicle and Telogator2 to estimate the telomere length from the long read sequences of each cancer cell line and compared it to the Telo-seq estimates. Results from both methods were highly correlated with the telomere length from Telo-seq (Fig 8A), and Topsicle had higher correlations for 9 of the cancer cell line dataset. Topsicle also had lower memory usage and computing time compared to Telogator2 (Fig 8B and 8C).

**Figure 8.**
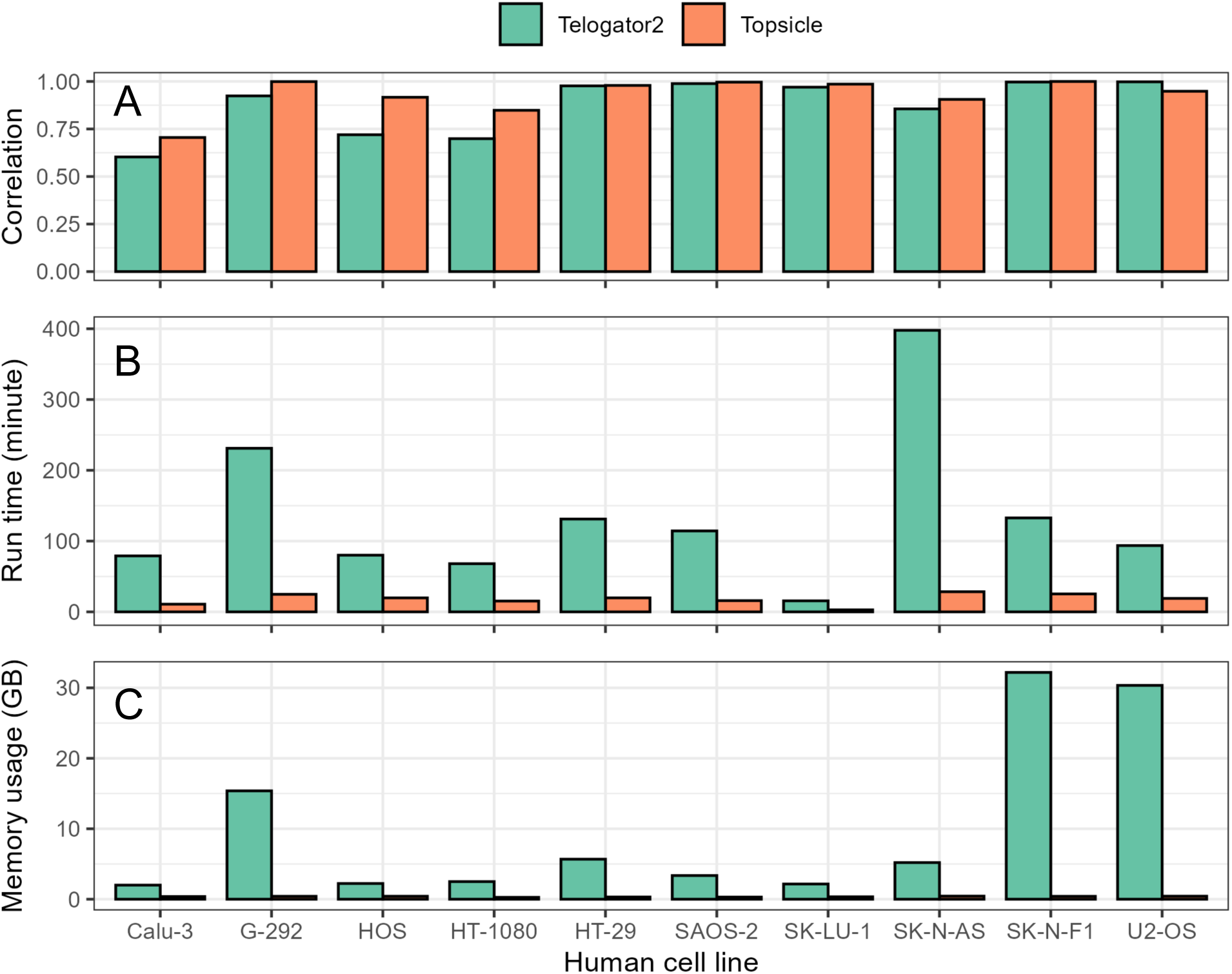
Comparing the performance of Topsicle and Telogator2o n long read sequences from human cancer cell lines. (A) Pearson’s correlation coefficient calculated by comparing telomere length estimates from Telo-seq and Telogator2 (green bars) or Topsicle (orange bars). Comparison of Telogator2 and Topsicle (B) run time and (C) memory usage.

## Discussion

The biological significance of telomere length variation has been a great interest to biomedical [99] and eco-evolutionary studies [100,101], but a largely open question is why the telomere length varies between species or between individuals of the same species and what is its function. A major step towards addressing this question involves measuring the length of the telomere in a wide range of organisms and through comparative analysis of the length variation with varying life history strategies and traits [102,103]. Here we introduce Topsicle, a new computational method that takes advantage of the reads generated from long read sequencing platforms, and estimates accurate and high-resolution telomere lengths from the sequencing data. The major advantages of our method is that it does not require a chromosome-level reference genome assembly or an experimental step to concentrate telomere long reads, and it can be applied to any organism with a telomerase based mechanism of extending short telomere repeat sequences at chromosome ends. Our method does require a prior knowledge of the telomere repeat sequence that will be searched across the sequencing library, but this can be determined from telomere sequence databases [104,105] or the repeat motif can be identified de novo using tandem repeat detection methodologies [33,106,107]. Overall, we predict Topsicle would be a useful computational method for estimating the length of the telomere for many organisms and assist telomere biology studies that are aimed at understanding the biological significance of the length variation.

Topsicle can search the entire sequencing library to determine reads sequenced from the telomere–subtelomere region, which greatly increases the accessibility of estimating telomere length to whole genome sequencing data without any specialized requirements. Topsicle calculates TRC values for all long reads to search for high telomere repeat content in the start or end of the reads and use those reads that pass a user defined TRC cutoff to calculate telomere length. Through this process it’s possible to include interstitial telomeric sequences [108,109], which would result in false telomere length estimates. We recommend using a series of stringent TRC cutoff and k-mer sizes to investigate the changes in the telomere length estimates, for example as we discovered in maize B73 a less stringent parameter for Topsicle resulted in an over estimation of telomere lengths. Topsicle is also able to analyze the telomeric regions of genome assemblies and complement the estimations made from raw long read sequences. Though the accuracy of assembling a repetitive region such as the telomere is not clear and caution the analysis of genome assembly based length measurements.

By using a change point detection method Topsicle utilizes the underlying repeat density to detect rapid changes in telomere repeat content to estimate the telomere length. This sharp transition of telomere repeat to non-telomere sequence has been a commonly used metric for determining the end of a telomere [52,57,97], and we use the signal to demarcate the telomere– subtelomere boundary for telomere length estimation. We applied Topsicle on three example plant species that greatly differed in its genome size and the overall repeat content (*i.e. A. thaliana* and maize) or had naturally varying telomere repeat sequences (*i.e. Mimulus* species group). Telomere length estimated from Topsicle were highly correlated with the gold standard TRF based estimates, and in the case of *Mimulus* the Topsicle based estimates were nearly identical to the TRF based telomere length estimates. On the other hand, for *A. thaliana* and maize the lengths from Topsicle were often shorter (∼1 kbp for *A. thaliana* and maize) compared to TRF based telomere length. Similar results have also been observed from long read sequences of human telomeres where computational estimates were several thousand basepairs shorter compared to TRF estimates [110]. This can be partially explained by the methodology of TRF where the restriction enzyme cut sites are aimed to digest the subtelomere region, and the extra subtelomere sequences will be included in the length estimates. It should be noted the quality of the library preparation would affect the length estimate and this could also explain the length discrepancy between Topsicle and TRF. Although from our maize analysis, there were no significant differences between libraries sequenced from the same genotype, suggesting minimal variation in length measurements at least between sequencing libraries. A future study testing library preparation methods and the resulting Topsicle estimated telomere lengths, could clarify our understanding of how much of the estimated telomere length variation arises from differences in generating the sequencing library.

Our analysis discovered the reads from the Nanopore sequencing platform had unique errors involving the telomere repeats that were not observed in the PacBio sequencing reads. Basecalling errors in Nanopore telomere reads have been detected in several organisms, including *A. thaliana* where CCCTAAA were miscalled as CCTGGG [73]. With our k-mer analysis we also discovered CCCTAAA was miscalled as CCCTAGG (Fig 1B) and we found additional errors involving the homopolymer repeat sequence. These errors can be fixed by re-basecalling the Nanopore data with a basecaller trained from a telomere model [73], but this is not always possible since a high-quality ground truth sequencing reads from the telomere are required for the training and the raw Nanopore sequencing signal data may not be available. Topsicle can overcome these sequencing errors by using smaller sized telomere repeat k-mers at different phases and count the telomere repeat sequence that is unaffected by the basecalling error. For instance, our analysis showed that by using a 4-mer telomere repeat, Topsicle estimated telomere length with the Nanopore sequencing data were not significantly different from the PacBio sequencing data for *A. thaliana* Col-0. But when using a perfect matching 7-mer telomere repeat sequence on the Nanopore reads, the telomere length estimates were significantly lower compared to the 4-mer telomere repeat based telomere length estimates. Our simulations showed that increasing sequencing errors resulted in a reduction of telomere length estimates, suggesting telomere length estimates from Nanopore reads would be underestimated if the errors are not taken into account. In addition, naturally varying sequence heterogeneous telomeres are commonly found in eukaryotic telomeres [64,100] and this would complicate computational methods estimating telomere lengths. For instance, from our analysis we discovered in *M. lewisii*, which has a sequence heterogeneous telomere consisting of both AAACCG and AAACCCG telomere repeats [68], it’s telomere length were underestimated when a exact matching AAACCG 6-mer repeat were used for length estimation. However, applying the strategy implemented in Topsicle and using a shorter telomere k-mer repeat the Topsicle based telomere length estimates were almost identical to TRF based telomere length estimates. This suggests using different sized k-mer repeats is a promising approach to estimate length of telomeres in various scenarios.

## Conclusion

In summary, Topsicle can accurately estimate telomere lengths from long read sequences. Importantly it adds to the available telomere length measuring toolkits, especially by allowing the analysis of whole genome long read sequencing datasets with minimal requirements. We hope Topsicle would assist the computational analysis of telomere length in various organisms and facilitate our understanding of the telomere length variation.

## Methods

### Topsicle algorithm

The overall strategy of Topsicle involves 1) identifying telomere repeat k-mers in a sequencing read, 2) calculating density of telomere repeats, and 3) using the density to estimate telomere length by applying a change point detection method. Topsicle accepts FASTQ/FASTA long read sequences and zipped versions of them as the input data and these reads can be those that are aligned to chromosome ends (*i.e.* from telomere and subtelomere region) or all whole genome sequencing reads. Initially, a basic quality control process is implemented where for each read will be checked for a minimum length threshold (parameter - minSeqLength) and the initial sequences of the reads can be trimmed on both ends of the reads (designated by parameter -trimfirst) to remove potential adapter sequences.

Topsicle will then search for the user defined telomere repeat sequence (parameter -pattern) and use the user defined k-mer size (parameter -telophrase) to search for telomere repeat patterns in all input reads. Reads that pass the TRC value cutoff (decided by user with parameter -cutoff) will be used for telomere length estimation. TRC value is measured for all input reads by checking the first or last 1,000 bp of a sequencing read and the side with the higher TRC value is chosen. TRC is calculated as the proportion of observed telomere repeat given the expected repeat count. For example, for the 6 basepair telomere repeat AAACCG a 1,000 bp would expect to hold 166 repeats and with a TRC cutoff of 0.4 it will only analyze reads with at least 66 repeats in the first or last 1,000 bp of a sequencing read. Reads that pass the TRC cutoff were subjected to a sliding window analysis (parameter -windowSize determines size of the window and -slide determines the increment size), where in each window it counts the total number of k-mer repeat for all phase of the k-mer sequence. The average k-mer counts are then calculated for the window. To determine the telomere–subtelomere boundary, the k-mer count windows are scanned and change point is detected using the binary segmentation algorithm [83] from the ruptures package [111]. A user defined maximum telomere length filter can be implemented (parameter - maxlengthtelo) to remove potential interstitial telomere repeats (e.g. maximum telomere length was set as 20 kbp for *A. thaliana* and *Mimulus*, while as 60 kbp for maize). Topsicle is written in Python3 and is available at https://github.com/jaeyoungchoilab/Topsicle.

### Simulation analysis

Performance of Topsicle was assessed through simulations that generated telomere reads with varying length and error rates. We constructed an artificial sequence representative of the telomere–subtelomere region by repeating the *A. thaliana* telomere repeat sequence AAACCCT multiple times and the remainder of the sequence filled with random nucleotides. Variable proportions of the telomere repeat were used for simulating long read sequences. We used PBSIM3 [112] on the artificial telomere–subtelomere region as a reference to simulate Nanopore long reads. Error rates were incorporated using the ERRHMM-ONT model with parameter -accuracy-mean to adjust the overall accuracy rate of sequencing to 0.7, 0.8 and 0.9. Reads are simulated 30 times and tested their telomere lengths with Topsicle.

### Asymptotic TRC threshold calculation

For automatically deciding an appropriate TRC threshold we implemented a asymptotic method of threshold determination. For each sequencing we calculated the telomere length and its TRC value. A quadratic function was then fit using the equation:

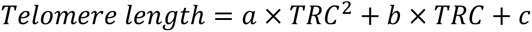

To calculate the asymptotic TRC threshold we calculated the derivative of the quadratic equation set to zero.

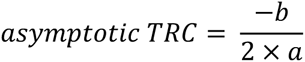

### A. thaliana data analysis

Long read sequences from the two prior studies [85,89] were downloaded from the Sequence Read Archive (SRA). For chromosome-end specific analysis, reads were aligned to the *A. thaliana* Col-0 chromosome-level genome assembly using minimap2 [113] with options -L -ax map-ont for ONT sequences and -L -ax map-pb for PacBio sequences. Sequencing reads from chromosome ends were extracted from the sorted genome alignment BAM file using seqkit [114] and selecting for reads found in the first and last 4000 bp of each chromosome. We used Topsicle with parameters of a TRC cutoff of 0.4 and using the 4-mer repeat (based on AAACCCT) to estimate the telomere lengths from the whole genome sequencing data.

### Maize data analysis

Long read sequences of 27 maize genotypes [48,95] were downloaded from Sequence Read Archive (SRA), then analyzed using Topsicle with the 5-mer repeat sequence (based on AAACCCT) and a TRC cutoff of 0.4.

### Mimulus data analysis

Mimulus reference genomes were downloaded from Mimubase (http://mimubase.org/). The genome versions of each species we used are as follows: M. cardinalis v2.0 (CE10g_v2.0), M. lewisii v2.0 (LF10g_v2.0), and M. verbenaceus v2.0 (MvBLg_v2.0). The whole genome sequencing Nanopore data were downloaded from SRA under the identifier PRJNA1049363. For chromosome-end specific telomere length analysis, each species was aligned to its reference genome with similar procedures as in *A. thaliana* data analysis, and we used Topsicle parameters of the 4-mer repeat sequence (based on AAACCG) and a TRC cutoff of 0.4.

### Human cancer cell line analysis

The long read sequences from Telo-seq study [58] was obtained from SRA under the identifier PRJNA1040425. For each cell line, we randomly extracted 2000 reads that were reported to have telomere sequences from Schmidt et al. [58] and ran Telogator2 with option -r ont and -n 4; and we ran Topsicle with parameters TRC cut off of 0.2, minimum read length is 3,000 bp and maximum read length is 50,000 bp. Computing time and memory usage were measured using the seff command available on the slurm environment.

## Declarations

### Ethics approval and consent to participate

Not applicable.

### Consent for publication

Not applicable.

### Availability of data and materials

Data from previously published study used for our analysis were obtained from Sequence Read Archive under accession numbers PRJCA012695, PRJEB31061, PRJEB32225, PRJEB62038, PRJNA751841, PRJNA1040425, and PRJNA1049363. Topsicle program is available on GitHub (https://github.com/jaeyoungchoilab/Topsicle) and the version used in the manuscript is available from Zenodo (10.5281/zenodo.15576160).

### Authors’ contributions

J.Y.C was responsible for the supervision and administration of the project. L.N. and J.Y.C developed Topsicle and the underlying methodology. L.N. performed all analysis. L.N. and J.Y.C wrote the manuscript and prepared all figures and tables. All authors read and approved the manuscript.

## Acknowledgements

We thank the weekly evolutionary genomics lab meeting members from the Choi lab members (Surbhi Kumawat, Naseem Samo, and Askhan Shametov), Katya Mack lab members (Brandon Meter, Vinay Sagar, and Miles Whedbee), and Austin Nguyen for valuable feedback on our research. We also thank Kevin Wei and Jamie Walters for their advice on the data analysis.

## Additional File 1: Fig S1-S12

Figure S1. Distribution of Telomere Repeat Count (TRC) values from Nanopore sequencing long reads. In each panel TRC values were calculated using 4-mer, 5-mer, 6-mer, or 7-mer telomere repeat for the *A. thaliana* Col-0 Nanopore reads that were aligning to chromosome ends and visually categorized as Type I, II, or III reads.

Figure S2. Telomere length estimates from the Nanopore sequencing or PacBio sequencing read data. Red stars indicate significant difference (p-value <0.05) after Mann-Whitney U test.

Figure S3. Distribution of Telomere Repeat Count (TRC) values from PacBio sequencing long reads. In each panel TRC values were calculated using 6-mer or 7-mer telomere repeat for the maize B73 PacBio reads that align to chromosome end or all reads from the genome.

Figure S4. Applying Topsicle on simulated dataset. Telomere length was estimated with Topsicle using the 5-mer, 6-mer, or 7-mer telomere repeat and TRC value cutoff of 0.4 on 30 simulated reads with error rates of 10%, 20%, and 30% for reads with varying length and proportion of telomere repeat. For each simulation the read length is indicated on the right side of the bar (“|”) and the length of the telomere repeat is on the left side of the bar.

Figure S5. Applying Topsicle on simulated dataset with low sequencing errors. Telomere length was estimated with Topsicle using the 4-mer or 7-mer telomere repeat and TRC value cutoff of 0.4 on 30 simulated reads with error rates of 1%, 2%, and 5% for reads with varying length and proportion of telomere repeat. For each simulation the read length is indicated on the right side of the bar (“|”) and the length of the telomere repeat is on the left side of the bar.

Figure S6. Scatter plot of telomere length estimates from 104 *A. thaliana* ecotype long read sequencing data. Telomere lengths were estimated using reads aligning to chromosome ends or from the whole genome sequencing data. Topsicle was used for length estimation with 4-mer telomere repeat and a TRC value cutoff of 0.4. Pearson’s r = 0.93 and p-value < 0.0001; Spearman’s ρ = 0.94 and p-value < 0.0001.

Figure S7. Counts of the reads that were used by Topsicle to estimate telomere length. Each row corresponds to a long read sequencing data of an *A. thaliana* ecotype from the Kang et al. (2023) or Lian et al. (2024) study. Each cell is the number of reads used by Topsicle for length estimation using reads that were aligning to chromosome ends, reads that were selected from the entire whole genome sequencing data, or reads that were found in both cases.

Figure S8. Scatter plot of the median Topsicle based telomere length of the 104 *A. thaliana* ecotypes and its genome sequencing statistics.

Figure S9. Distribution of the difference in telomere length estimated by TRF and Topsicle for the 31 *A. thaliana* ecotypes.

Figure S10. Scatter plot of 31 *A. thaliana* ecotypes with Topsicle based telomere length estimation and k-Seek based telomere repeat abundance.

Figure S11. Correlation of telomere length estimates using the asymptotic TRC threshold or manual threshold (TRC = 0.4) TRC cutoff. *A. thaliana* long read sequencing data from Lian et al. (2024) were used for the analysis.

Figure S12. Boxplot of Topsicle based telomere length estimates for the three *Mimulus* species. Varying k-mer size (based on the AAACCG telomere repeat sequence) and TRC cutoffs were used for estimating the telomere length.

## Additional File 2: Table S1-S6

Table S1. Comparison of telomere length estimates from Topsicle analyzing genome assembly or raw long read sequences.

Table S2. Telomere length statistics for the 104 *A. thaliana* ecotypes with long read sequencing data.

Table S3. Proportion of *A. thaliana* ecotypes with a chromosome specific telomere length.

Table S4. Topsicle based telomere length estimates for the sequencing libraries of the 27 maize genotypes.

Table S5. ANOVA F-statistic table for the 27 maize genotypes testing variability in telomere length between sequencing libraries.

Table S6. *In silico* TRF analysis on maize B73 genotype.

## Reference

1. O’Sullivan RJ, Karlseder J. Telomeres: protecting chromosomes against genome instability. Nat Rev Mol Cell Biol. 2010;11:171–81.

2. Olovnikov AM. A theory of marginotomy: The incomplete copying of template margin in enzymic synthesis of polynucleotides and biological significance of the phenomenon. J Theor Biol. 1973;41:181–90.

3. Watson JD. Origin of Concatemeric T7DNA. Nature New Biol. 1972;239:197–201.

4. de Lange T. How Telomeres Solve the End-Protection Problem. Science. 2009;326:948.

5. Blackburn EH. Telomeres and telomerase: their mechanisms of action and the effects of altering their functions. FEBS Lett. 2005;579:859–62.

6. de Lange T. Shelterin-Mediated Telomere Protection. Annu Rev Genet. 2018;52:223–47.

7. Ferreira MG, Miller KM, Cooper JP. Indecent Exposure: When Telomeres Become Uncapped. Mol Cell. 2004;13:7–18.

8. McKnight TD, Shippen DE. Plant telomere biology. Plant Cell. 2004;16:794–803.

9. Shay JW, Wright WE. Senescence and immortalization: role of telomeres and telomerase. Carcinogenesis. 2005;26:867–74.

10. Stanley SE, Armanios M. The short and long telomere syndromes: paired paradigms for molecular medicine. Curr Opin Genet Dev. 2015;33:1–9.

11. Armanios M. The Role of Telomeres in Human Disease. Annu Rev Genomics Hum Genet. 2022;23:363–81.

12. Calado RT, Dumitriu B. Telomere Dynamics in Mice and Humans. Semin Hematol. 2013;50:165–74.

13. Samassekou O, Gadji M, Drouin R, Yan J. Sizing the ends: Normal length of human telomeres. Ann Anat - Anat Anz. 2010;192:284–91.

14. Choi JY, Abdulkina LR, Yin J, Chastukhina IB, Lovell JT, Agabekian IA, et al. Natural variation in plant telomere length is associated with flowering time. Plant Cell. 2021;33:1118–34.

15. Hayflick L, Moorhead PS. The serial cultivation of human diploid cell strains. Exp Cell Res. 1961;25:585–621.

16. van Deursen JM. The role of senescent cells in ageing. Nature. 2014;509:439–46.

17. Aubert G, Lansdorp PM. Telomeres and Aging. Physiol Rev. 2008;88:557–79.

18. Young AJ. The role of telomeres in the mechanisms and evolution of life-history trade-offs and ageing. Philos Trans R Soc B Biol Sci [Internet]. 2018 [cited 2020 Jun 28];373. Available from: https://www.ncbi.nlm.nih.gov/pmc/articles/PMC5784072/

19. Whittemore K, Vera E, Martínez-Nevado E, Sanpera C, Blasco MA. Telomere shortening rate predicts species life span. Proc Natl Acad Sci U S A. 2019;116:15122–7.

20. Watson JM, Riha K. Telomeres, aging, and plants: from weeds to Methuselah - a mini-review. Gerontology. 2011;57:129–36.

21. Campitelli BE, Razzaque S, Barbero B, Abdulkina LR, Hall MH, Shippen DE, et al. Plasticity, pleiotropy and fitness trade-offs in Arabidopsis genotypes with different telomere lengths. New Phytol. 2022;233:1939–52.

22. Harley CB, Futcher AB, Greider CW. Telomeres shorten during ageing of human fibroblasts. Nature. 1990;345:458–60.

23. Kimura M, Stone RC, Hunt SC, Skurnick J, Lu X, Cao X, et al. Measurement of telomere length by the Southern blot analysis of terminal restriction fragment lengths. Nat Protoc. 2010;5:1596–607.

24. Lai T-P, Wright WE, Shay JW. Generation of digoxigenin-incorporated probes to enhance DNA detection sensitivity. BioTechniques. 2016;60:306–9.

25. Lansdorp PM, Verwoerd NP, van de Rijke FM, Dragowska V, Little M-T, Dirks RW, et al. Heterogeneity in Telomere Length of Human Chromosomes. Hum Mol Genet. 1996;5:685–91.

26. Rufer N, Dragowska W, Thornbury G, Roosnek E, Lansdorp PM. Telomere length dynamics in human lymphocyte subpopulations measured by flow cytometry. Nat Biotechnol. 1998;16:743–7.

27. Cawthon RM. Telomere measurement by quantitative PCR. Nucleic Acids Res. 2002;30:e47.

28. Aviv A. Commentary: Raising the bar on telomere epidemiology. Int J Epidemiol. 2009;38:1735–6.

29. Baird DM, Rowson J, Wynford-Thomas D, Kipling D. Extensive allelic variation and ultrashort telomeres in senescent human cells. Nat Genet. 2003;33:203–7.

30. Heacock M, Spangler E, Riha K, Puizina J, Shippen DE. Molecular analysis of telomere fusions in Arabidopsis: multiple pathways for chromosome end-joining. EMBO J. 2004;23:2304– 13.

31. Aubert G, Hills M, Lansdorp PM. Telomere length measurement-caveats and a critical assessment of the available technologies and tools. Mutat Res. 2012;730:59–67.

32. Ding Z, Mangino M, Aviv A, UK10K Consortium, Spector T, Durbin R. Estimating telomere length from whole genome sequence data. Nucleic Acids Res. 2014;42:e75.

33. Wei KH-C, Grenier JK, Barbash DA, Clark AG. Correlated variation and population differentiation in satellite DNA abundance among lines of Drosophila melanogaster. Proc Natl Acad Sci U S A. 2014;111:18793–8.

34. Wei KH-C, Lower SE, Caldas IV, Sless TJS, Barbash DA, Clark AG. Variable Rates of Simple Satellite Gains across the Drosophila Phylogeny. Mol Biol Evol. 2018;35:925–41.

35. Nersisyan L, Arakelyan A. Computel: Computation of Mean Telomere Length from Whole-Genome Next-Generation Sequencing Data. PLOS ONE. 2015;10:e0125201.

36. Farmery JHR, Smith ML, Lynch AG. Telomerecat: A ploidy-agnostic method for estimating telomere length from whole genome sequencing data. Sci Rep. 2018;8:1300.

37. Feuerbach L, Sieverling L, Deeg KI, Ginsbach P, Hutter B, Buchhalter I, et al. TelomereHunter – in silico estimation of telomere content and composition from cancer genomes. BMC Bioinformatics. 2019;20:272.

38. Yu Y, Mai Y, Zheng Y, Shi L. Assessing and mitigating batch effects in large-scale omics studies. Genome Biol. 2024;25:254.

39. Adey A, Morrison HG, Asan, Xun X, Kitzman JO, Turner EH, et al. Rapid, low-input, low-bias construction of shotgun fragment libraries by high-density in vitro transposition. Genome Biol. 2010;11:R119.

40. Aird D, Ross MG, Chen W-S, Danielsson M, Fennell T, Russ C, et al. Analyzing and minimizing PCR amplification bias in Illumina sequencing libraries. Genome Biol. 2011;12:R18.

41. Benjamini Y, Speed TP. Summarizing and correcting the GC content bias in high-throughput sequencing. Nucleic Acids Res. 2012;40:e72–e72.

42. Zavala-Paez M, Holliday J, Hamilton JA. Leveraging whole-genome sequencing to estimate telomere length in plants. Mol Ecol Resour. 2024;24:e13899.

43. Marx V. Method of the year: long-read sequencing. Nat Methods. 2023;20:6–11.

44. Warburton PE, Sebra RP. Long-Read DNA Sequencing: Recent Advances and Remaining Challenges. Annu Rev Genomics Hum Genet. 2023;24:109–32.

45. Garg V, Bohra A, Mascher M, Spannagl M, Xu X, Bevan MW, et al. Unlocking plant genetics with telomere-to-telomere genome assemblies. Nat Genet. 2024;56:1788–99.

46. Li H, Durbin R. Genome assembly in the telomere-to-telomere era. Nat Rev Genet. 2024;25:658–70.

47. Miga KH, Koren S, Rhie A, Vollger MR, Gershman A, Bzikadze A, et al. Telomere-to-telomere assembly of a complete human X chromosome. Nature. 2020;585:79–84.

48. Chen J, Wang Z, Tan K, Huang W, Shi J, Li T, et al. A complete telomere-to-telomere assembly of the maize genome. Nat Genet. 2023;55:1221–31.

49. O’Donnell S, Yue J-X, Saada OA, Agier N, Caradec C, Cokelaer T, et al. Telomere-to-telomere assemblies of 142 strains characterize the genome structural landscape in Saccharomyces cerevisiae. Nat Genet. 2023;55:1390–9.

50. Zhou Y, Xiong J, Shu Z, Dong C, Gu T, Sun P, et al. The telomere-to-telomere genome of Fragaria vesca reveals the genomic evolution of Fragaria and the origin of cultivated octoploid strawberry. Hortic Res. 2023;10:uhad027.

51. Reed J, Kirkman LA, Kafsack BF, Mason CE, Deitsch KW. Telomere length dynamics in response to DNA damage in malaria parasites. iScience. 2021;24:102082.

52. Stephens Z, Ferrer A, Boardman L, Iyer RK, Kocher J-PA. Telogator: a method for reporting chromosome-specific telomere lengths from long reads. Bioinformatics. 2022;38:1788–93.

53. Colt K, Petrus S, Abramson BW, Mamerto A, Hartwick NT, Michael TP. Telomere Length in Plants Estimated with Long Read Sequencing [Internet]. bioRxiv; 2024 [cited 2025 Mar 2]. p. 2024.03.27.586973. Available from: https://www.biorxiv.org/content/10.1101/2024.03.27.586973v1

54. Brown MR, Manuel Gonzalez de La Rosa P, Blaxter M. tidk: a toolkit to rapidly identify telomeric repeats from genomic datasets. Bioinformatics. 2025;41:btaf049.

55. Sholes SL, Karimian K, Gershman A, Kelly TJ, Timp W, Greider CW. Chromosome-specific telomere lengths and the minimal functional telomere revealed by nanopore sequencing. Genome Res. 2022;32:616–28.

56. Tham C-Y, Poon L, Yan T, Koh JYP, Ramlee MK, Teoh VSI, et al. High-throughput telomere length measurement at nucleotide resolution using the PacBio high fidelity sequencing platform. Nat Commun. 2023;14:281.

57. Karimian K, Groot A, Huso V, Kahidi R, Tan K-T, Sholes S, et al. Human telomere length is chromosome end–specific and conserved across individuals. Science. 2024;384:533–9.

58. Schmidt TT, Tyer C, Rughani P, Haggblom C, Jones JR, Dai X, et al. High resolution long-read telomere sequencing reveals dynamic mechanisms in aging and cancer. Nat Commun. 2024;15:5149.

59. Kazda A, Zellinger B, Rössler M, Derboven E, Kusenda B, Riha K. Chromosome end protection by blunt-ended telomeres. Genes Dev. 2012;26:1703–13.

60. Nelson ADL, Shippen DE. Blunt-ended telomeres: an alternative ending to the replication and end protection stories. Genes Dev. 2012;26:1648–52.

61. Hotaling S, Kelley JL, Frandsen PB. Toward a genome sequence for every animal: Where are we now? Proc Natl Acad Sci U S A. 2021;118:e2109019118.

62. Marks RA, Hotaling S, Frandsen PB, VanBuren R. Representation and participation across 20 years of plant genome sequencing. Nat Plants. 2021;7:1571–8.

63. Meyne J, Ratliff RL, Moyzis RK. Conservation of the human telomere sequence (TTAGGG)n among vertebrates. Proc Natl Acad Sci U S A. 1989;86:7049–53.

64. Červenák F, Sepšiová R, Nosek J, Tomáška Ľ. Step-by-Step Evolution of Telomeres: Lessons from Yeasts. Genome Biol Evol. 2020;13:evaa268.

65. Kuznetsova V, Grozeva S, Gokhman V. Telomere structure in insects: A review. J Zool Syst Evol Res. 2020;58:127–58.

66. Peska V, Garcia S. Origin, Diversity, and Evolution of Telomere Sequences in Plants. Front Plant Sci [Internet]. 2020 [cited 2023 Jan 31];11. Available from: https://www.frontiersin.org/articles/10.3389/fpls.2020.00117

67. Lim J, Kim W, Kim J, Lee J. Telomeric repeat evolution in the phylum Nematoda revealed by high-quality genome assemblies and subtelomere structures. Genome Res. 2023;33:1947–57.

68. Kumawat S, Shametov A, Valeeva LR, Ju Y, Martinez I, Logeswaran D, et al. Monkeyflower (Mimulus) uncovers the evolutionary basis of the eukaryote telomere sequence variation. PLOS Genet. 2025;21:e1011738.

69. Závodník M, Fajkus P, Franek M, Kopecký D, Garcia S, Dodsworth S, et al. Telomerase RNA gene paralogs in plants – the usual pathway to unusual telomeres. New Phytol. 2023;239:2353–66.

70. Amarasinghe SL, Su S, Dong X, Zappia L, Ritchie ME, Gouil Q. Opportunities and challenges in long-read sequencing data analysis. Genome Biol. 2020;21:30.

71. Wenger AM, Peluso P, Rowell WJ, Chang P-C, Hall RJ, Concepcion GT, et al. Accurate circular consensus long-read sequencing improves variant detection and assembly of a human genome. Nat Biotechnol. 2019;37:1155–62.

72. Sereika M, Kirkegaard RH, Karst SM, Michaelsen TY, Sørensen EA, Wollenberg RD, et al. Oxford Nanopore R10.4 long-read sequencing enables the generation of near-finished bacterial genomes from pure cultures and metagenomes without short-read or reference polishing. Nat Methods. 2022;19:823–6.

73. Tan K-T, Slevin MK, Meyerson M, Li H. Identifying and correcting repeat-calling errors in nanopore sequencing of telomeres. Genome Biol. 2022;23:180.

74. Rhoads A, Au KF. PacBio Sequencing and Its Applications. Genomics Proteomics Bioinformatics. 2015;13:278–89.

75. Mitsuhashi S, Frith MC, Mizuguchi T, Miyatake S, Toyota T, Adachi H, et al. Tandem-genotypes: robust detection of tandem repeat expansions from long DNA reads. Genome Biol. 2019;20:58.

76. Delahaye C, Nicolas J. Sequencing DNA with nanopores: Troubles and biases. PLOS ONE. 2021;16:e0257521.

77. Nix DA, Courdy SJ, Boucher KM. Empirical methods for controlling false positives and estimating confidence in ChIP-Seq peaks. BMC Bioinformatics. 2008;9:523.

78. Schmid K, Yang Z. The Trouble with Sliding Windows and the Selective Pressure in BRCA1. PLOS ONE. 2008;3:e3746.

79. Mallick S, Gnerre S, Muller P, Reich D. The difficulty of avoiding false positives in genome scans for natural selection. Genome Res. 2009;19:922–33.

80. Fletcher W, Yang Z. The effect of insertions, deletions, and alignment errors on the branch-site test of positive selection. Mol Biol Evol. 2010;27:2257–67.

81. Jordan G, Goldman N. The effects of alignment error and alignment filtering on the sitewise detection of positive selection. Mol Biol Evol. 2012;29:1125–39.

82. Zheng Y, Lunetta KL, Liu C, Katrinli S, Smith AK, Miller MW, et al. An evaluation of the genome-wide false positive rates of common methods for identifying differentially methylated regions using illumina methylation arrays. Epigenetics. 2022;17:2241–58.

83. Bai J. Estimating Multiple Breaks One at a Time. Econom Theory. 1997;13:315–52.

84. Shakirov EV, Chen JJ-L, Shippen DE. Plant telomere biology: The green solution to the end-replication problem. Plant Cell. 2022;34:2492–504.

85. Lian Q, Huettel B, Walkemeier B, Mayjonade B, Lopez-Roques C, Gil L, et al. A pan-genome of 69 Arabidopsis thaliana accessions reveals a conserved genome structure throughout the global species range. Nat Genet. 2024;56:982–91.

86. Wang B, Yang X, Jia Y, Xu Y, Jia P, Dang N, et al. High-quality Arabidopsis thaliana Genome Assembly with Nanopore and HiFi Long Reads. Genomics Proteomics Bioinformatics. 2022;20:4–13.

87. Copenhaver GP, Pikaard CS. RFLP and physical mapping with an rDNA-specific endonuclease reveals that nucleolus organizer regions of Arabidopsis thaliana adjoin the telomeres on chromosomes 2 and 4. Plant J. 1996;9:259–72.

88. Richards EJ, Ausubel FM. Isolation of a higher eukaryotic telomere from Arabidopsis thaliana. Cell. 1988;53:127–36.

89. Kang M, Wu H, Liu H, Liu W, Zhu M, Han Y, et al. The pan-genome and local adaptation of Arabidopsis thaliana. Nat Commun. 2023;14:6259.

90. Truong C, Oudre L, Vayatis N. Selective review of offline change point detection methods. Signal Process. 2020;167:107299.

91. Barcenilla BB, Meyers AD, Castillo-González C, Young P, Min J-H, Song J, et al. Arabidopsis telomerase takes off by uncoupling enzyme activity from telomere length maintenance in space. Nat Commun. 2023;14:7854.

92. Burr B, Burr FA, Matz EC, Romero-Severson J. Pinning down loose ends: mapping telomeres and factors affecting their length. Plant Cell. 1992;4:953–60.

93. Schnable PS, Ware D, Fulton RS, Stein JC, Wei F, Pasternak S, et al. The B73 maize genome: complexity, diversity, and dynamics. Science. 2009;326:1112–5.

94. Jiao Y, Peluso P, Shi J, Liang T, Stitzer MC, Wang B, et al. Improved maize reference genome with single-molecule technologies. Nature. 2017;546:524–7.

95. Hufford MB, Seetharam AS, Woodhouse MR, Chougule KM, Ou S, Liu J, et al. De novo assembly, annotation, and comparative analysis of 26 diverse maize genomes. Science. 2021;373:655–62.

96. Brown AN, Lauter N, Vera DL, McLaughlin-Large KA, Steele TM, Fredette NC, et al. QTL Mapping and Candidate Gene Analysis of Telomere Length Control Factors in Maize (Zea mays L.). G3 GenesGenomesGenetics. 2011;1:437–50.

97. Tao Y, Xian W, Bao Z, Rabanal FA, Movilli A, Lanz C, et al. Atlas of telomeric repeat diversity in Arabidopsis thaliana. Genome Biol. 2024;25:244.

98. Stephens Z, Kocher J-P. Characterization of telomere variant repeats using long reads enables allele-specific telomere length estimation. BMC Bioinformatics. 2024;25:194.

99. Chakravarti D, LaBella KA, DePinho RA. Telomeres: history, health, and hallmarks of aging. Cell. 2021;184:306–22.

100. Kumawat S, Choi JY. No end in sight: Mysteries of the telomeric variation in plants. Am J Bot. 2023;110:e16244.

101. Monaghan P. Linking telomere dynamics to evolution, life history and environmental change: perspectives, predictions and problems. Biogerontology. 2024;25:301–11.

102. Gomes NMV, Ryder OA, Houck ML, Charter SJ, Walker W, Forsyth NR, et al. Comparative biology of mammalian telomeres: hypotheses on ancestral states and the roles of telomeres in longevity determination. Aging Cell. 2011;10:761–8.

103. Pepke ML, Eisenberg DTA. On the comparative biology of mammalian telomeres: Telomere length co-evolves with body mass, lifespan and cancer risk. Mol Ecol. 2022;31:6286– 96.

104. Podlevsky JD, Bley CJ, Omana RV, Qi X, Chen JJ-L. The Telomerase Database. Nucleic Acids Res. 2008;36:D339–43.

105. Lyčka M, Bubeník M, Závodník M, Peska V, Fajkus P, Demko M, et al. TeloBase: a community-curated database of telomere sequences across the tree of life. Nucleic Acids Res. 2024;52:D311–21.

106. Benson G. Tandem repeats finder: a program to analyze DNA sequences. Nucleic Acids Res. 1999;27:573–80.

107. Chung G, Piano F, Gunsalus KC. TeloSearchLR: an algorithm to detect novel telomere repeat motifs using long sequencing reads [Internet]. bioRxiv; 2024 [cited 2025 Mar 19]. p. 2024.10.29.617943. Available from: https://www.biorxiv.org/content/10.1101/2024.10.29.617943v1

108. Regad F, Lebas M, Lescure B. Interstitial Telomeric Repeats within the *Arabidopsis thaliana* Genome. J Mol Biol. 1994;239:163–9.

109. Lin KW, Yan J. Endings in the middle: Current knowledge of interstitial telomeric sequences. Mutat Res Mutat Res. 2008;658:95–110.

110. Sanchez SE, Gu Y, Wang Y, Golla A, Martin A, Shomali W, et al. Digital telomere measurement by long-read sequencing distinguishes healthy aging from disease. Nat Commun. 2024;15:5148.

111. Truong C, Oudre L, Vayatis N. ruptures: change point detection in Python [Internet]. arXiv; 2018 [cited 2025 Mar 18]. Available from: http://arxiv.org/abs/1801.00826

112. Ono Y, Hamada M, Asai K. PBSIM3: a simulator for all types of PacBio and ONT long reads. NAR Genomics Bioinforma. 2022;4:lqac092.

113. Li H. Minimap2: pairwise alignment for nucleotide sequences. Bioinformatics. 2018;34:3094–100.

114. Shen W, Le S, Li Y, Hu F. SeqKit: A Cross-Platform and Ultrafast Toolkit for FASTA/Q File Manipulation. PLOS ONE. 2016;11:e0163962.

