## Additional File 1 for "Topsicle: a method for estimating telomere length from whole genome long-read sequencing data"

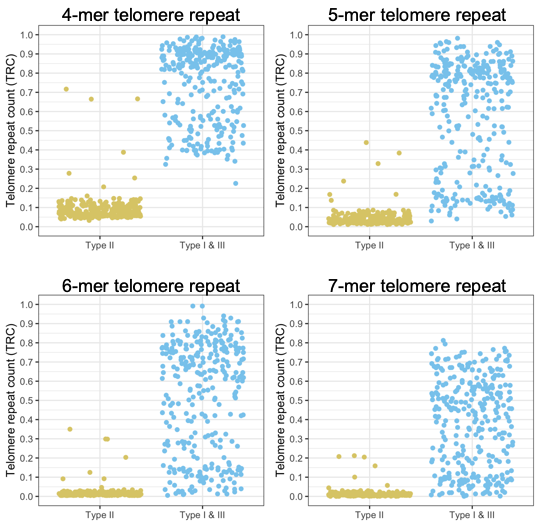

Figure S1. Distribution of Telomere Repeat Count (TRC) values from Nanopore sequencing long reads. In each panel TRC values were calculated using 4-mer, 5-mer, 6-mer, or 7-mer telomere repeat for the *A. thaliana* Col-0 Nanopore reads that were aligning to chromosome ends and visually categorized as Type I, II, or III reads.

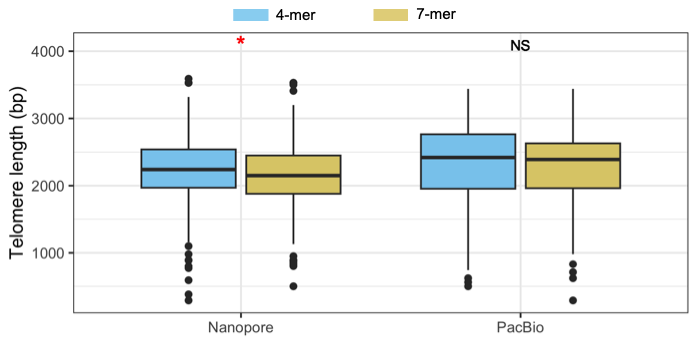

Figure S2. Telomere length estimates from the Nanopore sequencing or PacBio sequencing read data. Red stars indicate significant difference (p-value <0.05) after Mann-Whitney U test.

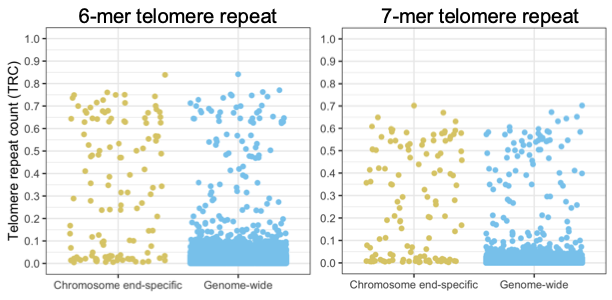

Figure S3. Distribution of Telomere Repeat Count (TRC) values from PacBio sequencing long reads. In each panel TRC values were calculated using 6-mer or 7-mer telomere repeat for the maize B73 PacBio reads that align to chromosome end or all reads from the genome.

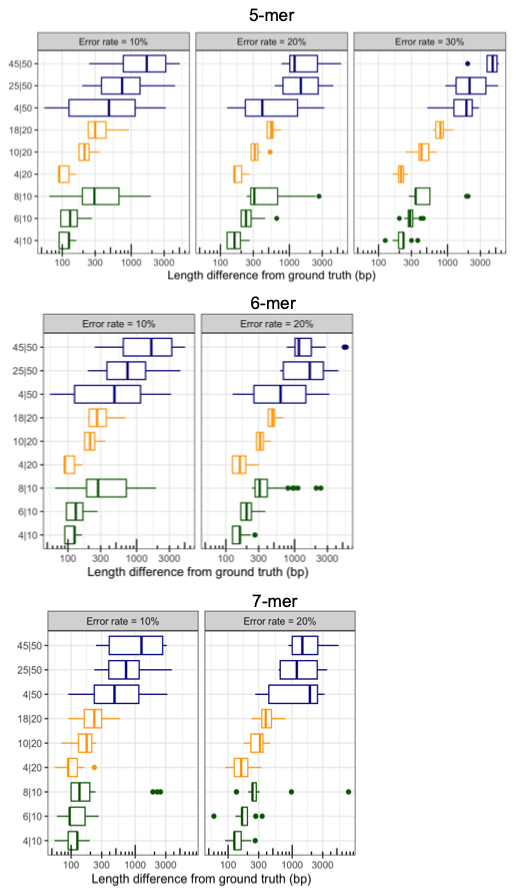

Figure S4. Applying Topsicle on simulated dataset. Telomere length was estimated with Topsicle using the 5-mer, 6-mer, or 7-mer telomere repeat and TRC value cutoff of 0.4 on 30 simulated reads with error rates of 10%, 20%, and 30% for reads with varying length and proportion of telomere repeat. For each simulation the read length is indicated on the right side of the bar (“|”) and the length of the telomere repeat is on the left side of the bar.

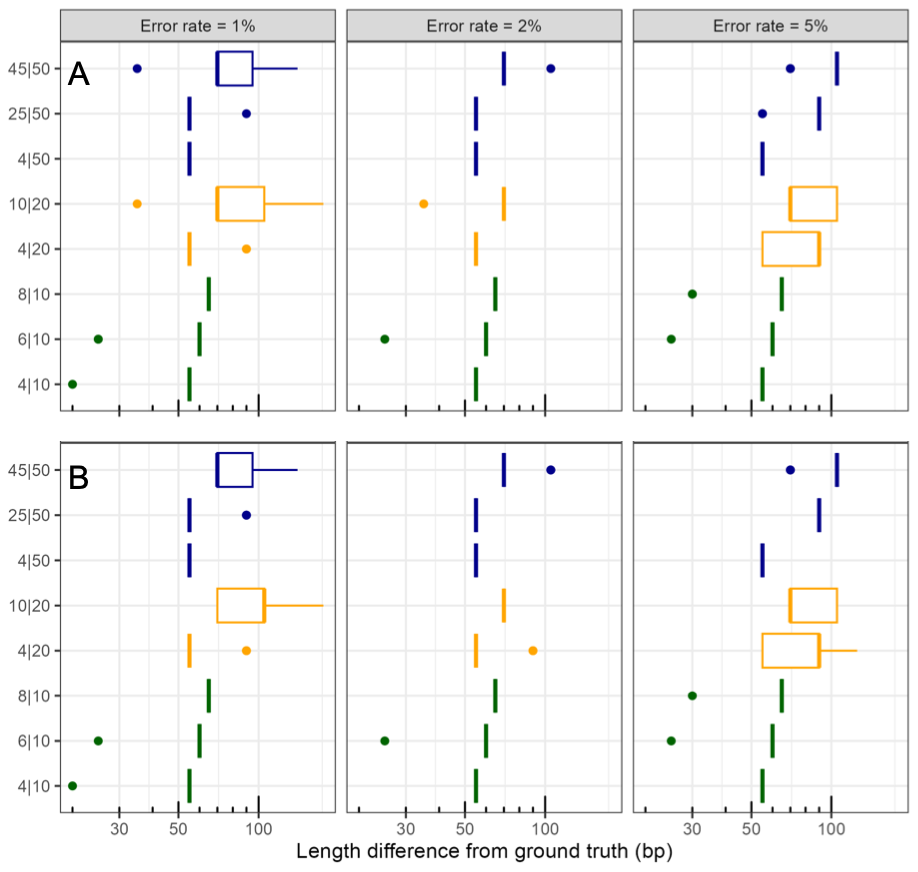

Figure S5. Applying Topsicle on simulated dataset with low sequencing errors. Telomere length was estimated with Topsicle using the 4-mer or 7-mer telomere repeat and TRC value cutoff of 0.4 on 30 simulated reads with error rates of 1%, 2%, and 5% for reads with varying length and proportion of telomere repeat. For each simulation the read length is indicated on the right side of the bar (“|”) and the length of the telomere repeat is on the left side of the bar.

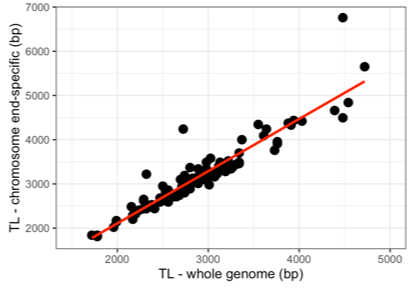

Figure S6. Scatter plot of telomere length estimates from 104 *A. thaliana* ecotype long read sequencing data. Telomere lengths were estimated using reads aligning to chromosome ends or from the whole genome sequencing data. Topsicle was used for length estimation with 4-mer telomere repeat and a TRC value cutoff of 0.4. Pearson’s r = 0.93 and p-value < 0.0001; Spearman’s ρ = 0.94 and p-value < 0.0001.

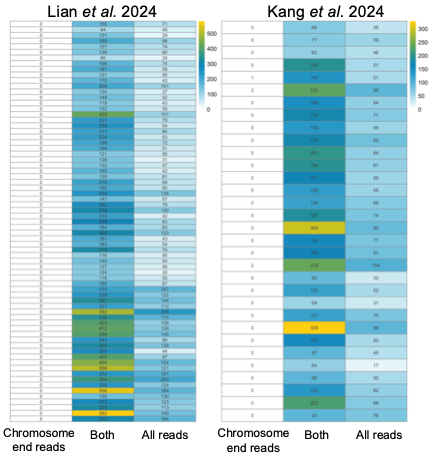

Figure S7. Counts of the reads that were used by Topsicle to estimate telomere length. Each row corresponds to a long read sequencing data of an *A. thaliana* ecotype from the Kang et al. (2023) or Lian et al. (2024) study. Each cell is the number of reads used by Topsicle for length estimation using reads that were aligning to chromosome ends, reads that were selected from the entire whole genome sequencing data, or reads that were found in both cases.

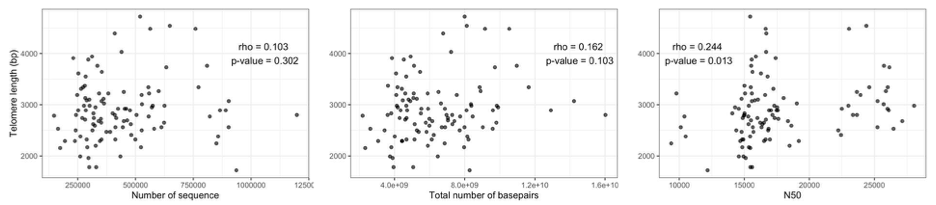

Figure S8. Scatter plot of the median Topsicle based telomere length of the 104 *A. thaliana* ecotypes and its genome sequencing statistics.

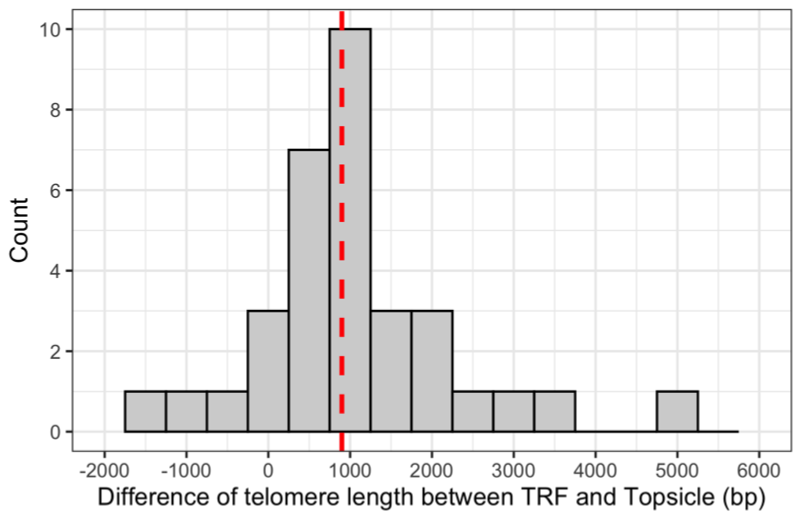

Figure S9. Distribution of the difference in telomere length estimated by TRF and Topsicle for the 31 *A. thaliana* ecotypes.

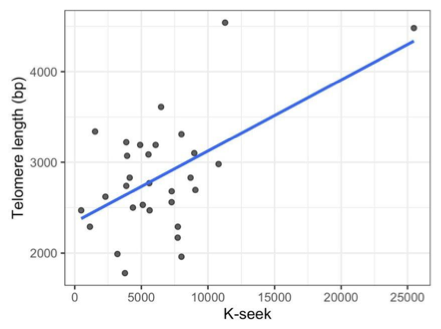

Figure S10. Scatter plot of 31 *A. thaliana* ecotypes with Topsicle based telomere length estimation and k-Seek based telomere repeat abundance.

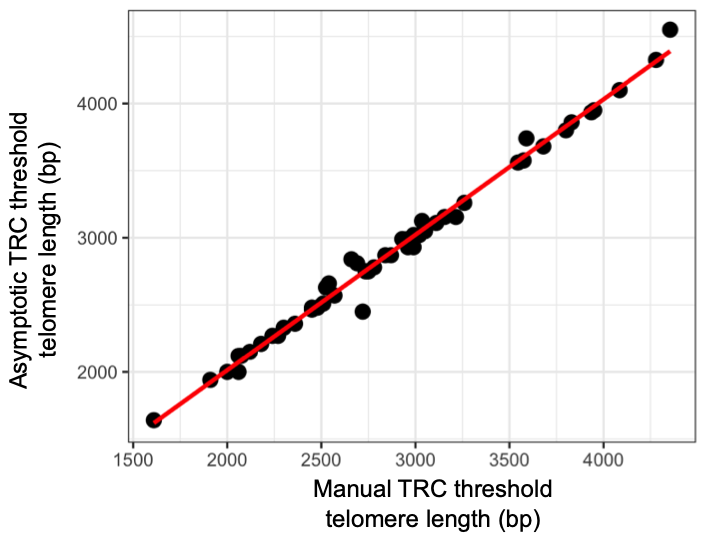

Figure S11. Correlation of telomere length estimates using the asymptotic TRC threshold or manual threshold (TRC = 0.4) TRC cutoff. *A. thaliana* long read sequencing data from Lian et al. (2024) were used for the analysis.

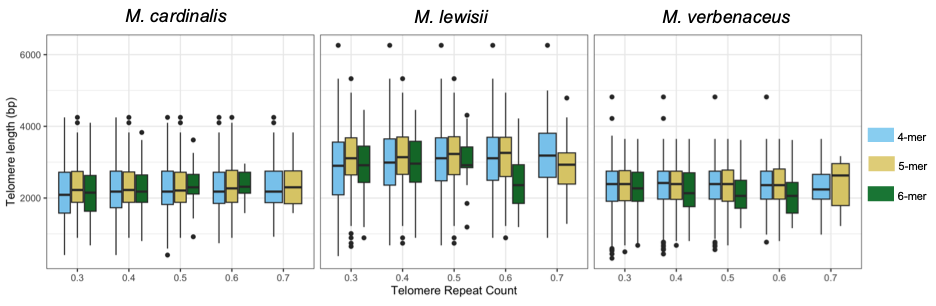
Figure S12. Boxplot of Topsicle based telomere length estimates for the three *Mimulus* species. Varying k-mer size (based on the AAACCG telomere repeat sequence) and TRC cutoffs were used for estimating the telomere length.
